# Refining the taxonomy of the order *Hyphomicrobiales* (*Rhizobiales*) based on whole genome comparisons of over 130 genus type strains

**DOI:** 10.1101/2023.11.15.567303

**Authors:** George C. diCenzo, Yuqi Yang, J. Peter W. Young, Nemanja Kuzmanović

**Author notes:** **Corresponding authors:** George diCenzo and Nemanja Kuzmanović (;).

## Abstract

The alphaproteobacterial order *Hyphomicrobiales* consists of 38 families comprising 155 validly published genera as of June 2023. The order *Hyphomicrobiales* was first described in 1957 and underwent important revisions in 2020. However, several inconsistencies in the taxonomy of this order remain, and there is a need for a consistent framework for defining families within the order. We propose a common genome-based framework for defining families within the order *Hyphomicrobiales*, suggesting that families represent monophyletic groups in core-genome phylogenies that share pairwise average amino acid identity values above ∼75% when calculated from a core set of 59 proteins. Applying this framework, we propose the formation of four new families and to reassign the genera *Salaquimonas*, *Rhodoblastus*, and *Rhodoligotrophos* into *Salaquimonadaceae* fam. nov., *Rhodoblastaceae* fam. nov., and *Rhodoligotrophaceae* fam. nov., respectively, and the genera *Albibacter*, *Chenggangzhangella*, *Hansschlegelia*, and *Methylopila* into *Methylopilaceae* fam. nov.. We further propose to unify the families *Bartonellaceae*, *Brucellaceae*, *Phyllobacteriaceae*, and *Notoacmeibacteraceae* as *Bartonellaceae*; the families *Segnochrobactraceae* and *Pseudoxanthobacteraceae* as *Segnochrobactraceae*; the families *Lichenihabitantaceae* and *Lichenibacteriaceae* as *Lichenihabitantaceae*; and the families *Breoghaniaceae* and *Stappiaceae* as *Stappiaceae*. Lastly, we propose to reassign several genera to existing families. Specifically, we propose to reassign the genus *Pseudohoeflea* to the family *Rhizobiaceae*; the genera *Oricola*, *Roseitalea*, and *Oceaniradius* to the family *Ahrensiaceae*; the genus *Limoniibacter* to the emended family *Bartonellaceae*; the genus *Faunimonas* to the family *Afifellaceae*; and the genus *Pseudochelatococcus* to the family *Chelatococcaceae*. Our data also support the recent proposal to reassign the genus *Prosthecomicrobium* to the family *Kaistiaceae*.

## INTRODUCTION

The order *Hyphomicrobiales* of the class *Alphaproteobacteria* consists of 38 families encompassing 155 valid genera (List of Prokaryotic names with Standing in Nomenclature; LPSN; accessed 25 June 2023). It was known by the name *Rhizobiales* Kuykendall 2006 [1] from 2005 until 2020, when Hördt and colleagues [2] pointed out that this was an illegitimate later synonym of *Hyphomicrobiales* Douglas 1957 (Approved List 1980) [3]. This order contains phenotypically diverse organisms, including terrestrial and aquatic bacteria, plant mutualists, and plant, animal, and human pathogens. Moreover, all known alpha-rhizobia (alphaproteobacterial nitrogen-fixing legume symbionts) are found within the order *Hyphomicrobiales* and are spread across seven families (*Rhizobiaceae*, *Phyllobacteriaceae*, *Brucellaceae*, *Nitrobacteriaceae*, *Methylobacteriaceae*, *Xanthobacteriaceae*, and *Devosiaceae*).

We recently proposed a genome-based framework for defining genera within the family *Rhizobiaceae* [4], which is the largest family within the order *Hyphomicrobiales* with 20 validly published genera as of 25 June 2023. While applying our framework to refine the family *Rhizobiaceae* [4], we wondered whether any genera currently assigned to the family *Rhizobiaceae* should be reassigned to novel families. This prompted us to evaluate family-level classifications within the order *Hyphomicrobiales*. Here, we propose a genome-based framework for defining families within the order *Hyphomicrobiales*. We then apply this framework and propose the unification of several families, the formation of four new families, and the reassignment of multiple genera.

## METHODS

### Datasets

Core-proteome phylogenies and overall genomic relatedness indices (OGRIs) were calculated using the genomes of 138 *Hyphomicrobiales* strains (**Dataset S1**). Where applicable, five *Caulobacterales* strains were included as an outgroup (**Dataset S2**). Most strains were genus type strains. The exceptions were the species type strains “*Pararhizobium mangrovi*” BGMRC 6574^T^, which was included as preliminary analyses suggested that this strain may belong to a new genus potentially in a novel family, and *Hyphomicrobium denitrificans* ATCC 51888^T^, which was included as the genome sequence of the *Hyphomicrobium* genus type strain was unavailable. Where noted, we included an additional four species type strains from the genus *Bartonella* and the recently proposed *“Flavimaribacter sediminis*” WL0058^T^ (**Dataset S3**). All genomes were downloaded from the National Center for Biotechnology Information (NCBI) Genome database.

For construction of a 16S rRNA gene phylogeny, full-length 16S rRNA gene sequences were extracted from the genome sequences of the 143 strains used in the genome-based analyses, where available. If the genome did not contain a full length 16S rRNA gene sequence, the strain’s 16S rRNA gene sequence was downloaded from NCBI GenBank via links embedded in the LPSN database [5]. In cases where more than one distinct 16S rRNA gene sequence was extracted from a whole genome sequence, all unique sequences were kept. This dataset was supplemented with the 16S rRNA gene sequences of nine genus type strains that lacked publicly available whole genome sequences but were related to strains that were candidates for reclassification. These 16S rRNA gene sequences were downloaded from the LPSN database (**Dataset S4**).

### Identification of core gene sets

Initially, the GET_HOMOLOGUES software package version 05052023 [6] and GET_PHYLOMARKERS software package version 2.4.5_17nov2022 [7] were used to identify non-recombining single-copy marker genes present in all 143 genomes, as described previously [8]. This led to the identification of 19 marker genes, which we termed “core_143”. Next, we used GET_HOMOLOGUES and custom scripts to identify single-copy marker genes present in at least 95% of the target genomes; the stringent filtering of GET_PHYLOMARKERS was not used. This led to the identification of 256 marker genes, which we termed “perc95_143”. During preliminary investigations of the perc95_143 gene set, we observed that five strains (*Chenggangzhangella methanolivorans* CHL1^T^, *Methylobrevis pamukkalensis* VKM B-2849^T^, *Nitratireductor aquibiodomus* JCM 21793^T^, *Liberibacter crescens* BT-1^T^, and *Methyloligella halotolerans* VKM B-2706^T^) lacked between 12% and 28% of these genes (see **Text S1**). We therefore repeated the above analyses using a reduced set of 138 strains that excluded those five strains. Two additional strains, *Segnochrobactrum spirostomi* Sp-1^T^ and *Ahrensia kielensis* DSM 5890^T^, lacked 9% and 7% of the perc95_143 genes, respectively; however, these two strains were kept in the dataset as they were the only representatives of their respective families. All other strains lacked less than 5% of the perc95_143 genes. Using the reduced dataset of 138 strains, we identified a core set of 59 non-recombining, single-copy marker genes present in all 138 strains (termed “core_138”) and 267 single-copy genes present in at least 95% of the target genomes (termed “perc95_138”). A summary of the four gene sets is provided in **Table 1**.

**Table 1.** Properties of the gene sets used for phylogenetic and overall genome similarity analyses.

|  | Gene set |  |  |  |
| --- | --- | --- | --- | --- |
|  | core_143 | perc95_143 | core_138 | perc95_138 |
| Number of *Hyphomicrobiales* strains | 138 | 138 | 133 | 133 |
| Number of *Caulobacterales* strains | 5 | 5 | 5 | 5 |
| Genomes containing the genes (%) | 100 | ≥95 | 100 | ≥95 |
| Number of genes | 19 | 256 | 59 | 267 |

### Calculation of overall genome relatedness indices

Core-proteome average amino acid identity (cpAAI) was computed as the proportion of differences (including gaps) in pairwise comparisons of protein sequences encoded by a given core gene set. These calculations were performed as described previously [4], and were dependent on the ‘ape’ package version 5.7-1 in R version 4.3.0 [9]. Whole-proteome average amino acid identity (wpAAI; usually simply known as AAI) was computed using EzAAI version 1.2.2 [10], with default parameters and the dependency Prodigal version 2.6.3 [11].

### Core-proteome phylogenies

For the core_143 and core_138 gene sets, GET_PHYLOMARKERS was used to align and concatenate the encoded proteins and to remove non-informative sites; alignment was performed using Clustal Omega version 1.2.4 [12]. For the perc95_143 and perc95_138 gene sets, the encoded proteins were aligned with MAFFT version 7.453 [13], after which the protein alignments were trimmed using trimAl version 1.4.rev22 [14] and the automated1 option, and then concatenated. For all four gene sets, the concatenated protein alignments were used as input for ModelFinder [15] as implemented in IQ-TREE version 2.2.2.4 [16], and the best scoring model was identified based on Bayesian information criterion (BIC). IQ-TREE was then used to create maximum-likelihood (ML) phylogenies from the concatenated alignments using the best-scoring model for each protein set (core_143: LG+F+R8; core_138: LG+F+R9; perc95_143: LG+F+I+R10; perc95_138: LG+F+I+R10). Branch supports were assessed in IQ-TREE using the Shimodaira-Hasegawa-like approximate likelihood ratio test (SH-aLRT) [17] and ultrafast jackknife analysis with a subsampling proportion of 40%, with both metrics calculated from 1000 replicates. Phylogenies were visualized using iTOL [18].

### 16S rRNA gene phylogenies

MAFFT was used to align the 16S rRNA gene sequences, after which trimAl was used to trim the alignment. We additionally aligned the 16S rRNA gene sequences using Clustal Omega with the --full and --full-iter options, and then trimmed the alignment with trimAl using the automated1 option. The trimmed nucleotide alignments were used as input for ModelFinder as implemented in IQ-TREE, which for both alignments, identified GTR+F+I+R6 as the best-scoring model based on BIC. IQ-TREE was then used to create ML phylogenies from both trimmed alignments, using the GTR+F+I+R6 model. Branch supports were assessed in IQ-TREE using SH-aLRT and an ultrafast bootstrap analysis, with both metrics calculated from 1000 replicates.

### Data availability

All genome sequences used in this work were previously published, and the assembly accessions are provided in **Datasets S1-S3**. Likewise, all 16S rRNA gene sequences used in this study were previously published, and the corresponding GenBank accessions for those not extracted directly from the whole genome sequences are provided in **Dataset S4**. Twelve supplementary figures, four supplementary datasets, and supplementary text are included in the online version of this article. All raw data (cpAAI data, wpAAI data, and Newick formatted phylogenies) used to generate the figures presented in this manuscript are available through FigShare (figshare.com/articles/online_resource/diCenzo_et_al_2023_Hyphomicrobiales_taxonomy/24417334). All code to repeat the analyses in this study is available through GitHub (github.com/diCenzo-Lab/012_2023_Hyphomicrobiales_taxonomy). Protein marker sets to allow for the identification of the marker proteins in other organisms using the cpAAI_Rhizobiaceae pipeline (github.com/flass/cpAAI_Rhizobiaceae) [4] are available through GitHub (github.com/diCenzo-Lab/012_2023_Hyphomicrobiales_taxonomy).

## RESULTS AND DISCUSSION

### Genome-based metrics for family-level delimitation in the order *Hyphomicrobiales*

To evaluate family assignments in the order *Hyphomicrobiales*, we began with a dataset of 138 *Hyphomicrobiales* strains, including 136 genus type strains, including at least one representative from each of the 38 families. Five *Caulobacterales* genus type strains were included as an outgroup. As in our recent study of the family *Rhizobiaceae* [4], we reasoned that genome sequence-based family delineation should consider both phylogenetic relatedness and genome similarity based on one or more overall genome relatedness indices (OGRIs). We chose to work with cpAAI and wpAAI as these appeared to be the most appropriate OGRIs based on our previous work [4]. Therefore, we first identified a set of 19 marker genes present in 100% of the 143 strains (termed core_143) and a set of 256 marker genes present in ≥95% of the strains (termed perc95_143). Both datasets were used to construct ML phylogenies (**Figures S1 and S2**), and core_143 was additionally used to calculate pairwise cpAAI values between all 143 strains (**Figure S1**, **Dataset S5**). Pairwise wpAAI was also calculated between all 143 strains (**Figures S3**, **Dataset S6**). Early analysis of these datasets revealed that five strains lacked >10% of the perc95_143 genes and that removal of these five strains significantly increased the size of the core genome (see **Text S1**). We therefore repeated all analyses using a dataset lacking these five strains. Using this reduced dataset, we identified a set of 59 marker genes present in 100% of the strains (termed core_138) and a set of 267 marker genes present in ≥95% of the strains (termed perc95_138). ML phylogenies were constructed from both core_138 (**Figure S4**) and perc95_138 (**Figure 1**), and pairwise cpAAI values were calculated between all 138 strains using core_138 (**Figure 1, Dataset S7**). A summary of the four gene sets is provided in **Table 1**.

**Figure 1.**
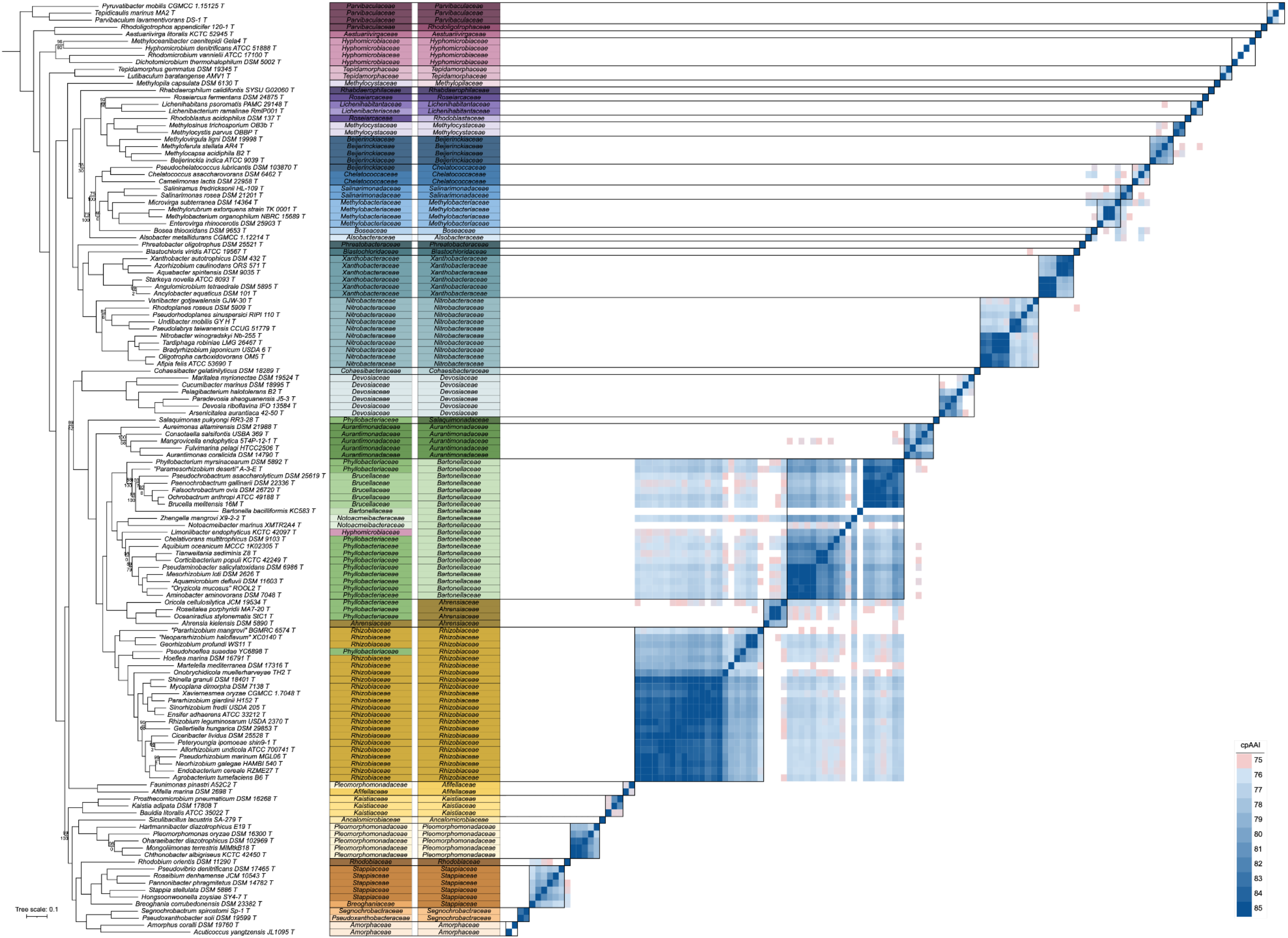
Phylogenetic and core-proteome AAI (cpAA) analyses of the order *Hyphomicrobiales*. On the left, a maximum likelihood phylogeny of 133 *Hyphomicrobiales* type strains is shown, built using the concatenated protein alignments encoded by the perc95_138 gene set (267 genes present in at least 95% of the strains). The phylogeny was rooted using five *Caulobacterales* type strains as the outgroup. The numbers on the nodes indicate the ultra-fast jackknife values using a 40% resampling rate (top numbers) and the SH-aLRT support values (bottom numbers), both calculated from 1000 replicates. Values are only shown at nodes where at least one value is below 100. The scale bar represents the average number of amino acid substitutions per site. To the right of the phylogeny is the current family assignment of each of the 133 *Hyphomicrobiales* type strains, followed to the right by the proposed family assignment of each strain. On the righthand side, a matrix is provided showing the cpAAI values between each pair of strains calculated using the proteins encoded by the core_138 gene set (59 genes present in 100% of the strains). Values less than 75% are in white while all values greater than 85% are the same shade of blue. Black boxes indicate the proposed families.

We wondered whether we could identify biologically relevant cpAAI or wpAAI thresholds for delineating families in the order *Hyphomicrobiales*. We therefore plotted the distribution of cpAAI values (calculated from core_138) as a histogram, plotting separate distributions for within-family, between-family but within-order, and between-order comparisons (**Figures 2A** and **2C**). In addition to using all cpAAI values calculated from core_138, we plotted the results for a reduced dataset lacking the family *Rhizobiaceae* (**Figure 2B** and **2D**). This was done for two reasons. The family *Rhizobiaceae* was over-represented in our dataset, accounting for over 15% of the strains in our analysis, and thus its inclusion had the potential to bias the distribution. In addition, the family *Rhizobiaceae* was not well separated from the related *Bartonellaceae* – *Phyllobacteriaceae* – *Brucellaceae* - *Notoacmeibacteraceae* clade (as discussed below), and thus had the potential to mask patterns that might otherwise be detected across the remaining 33 *Hyphomicrobiales* families.

**Figure 2.**
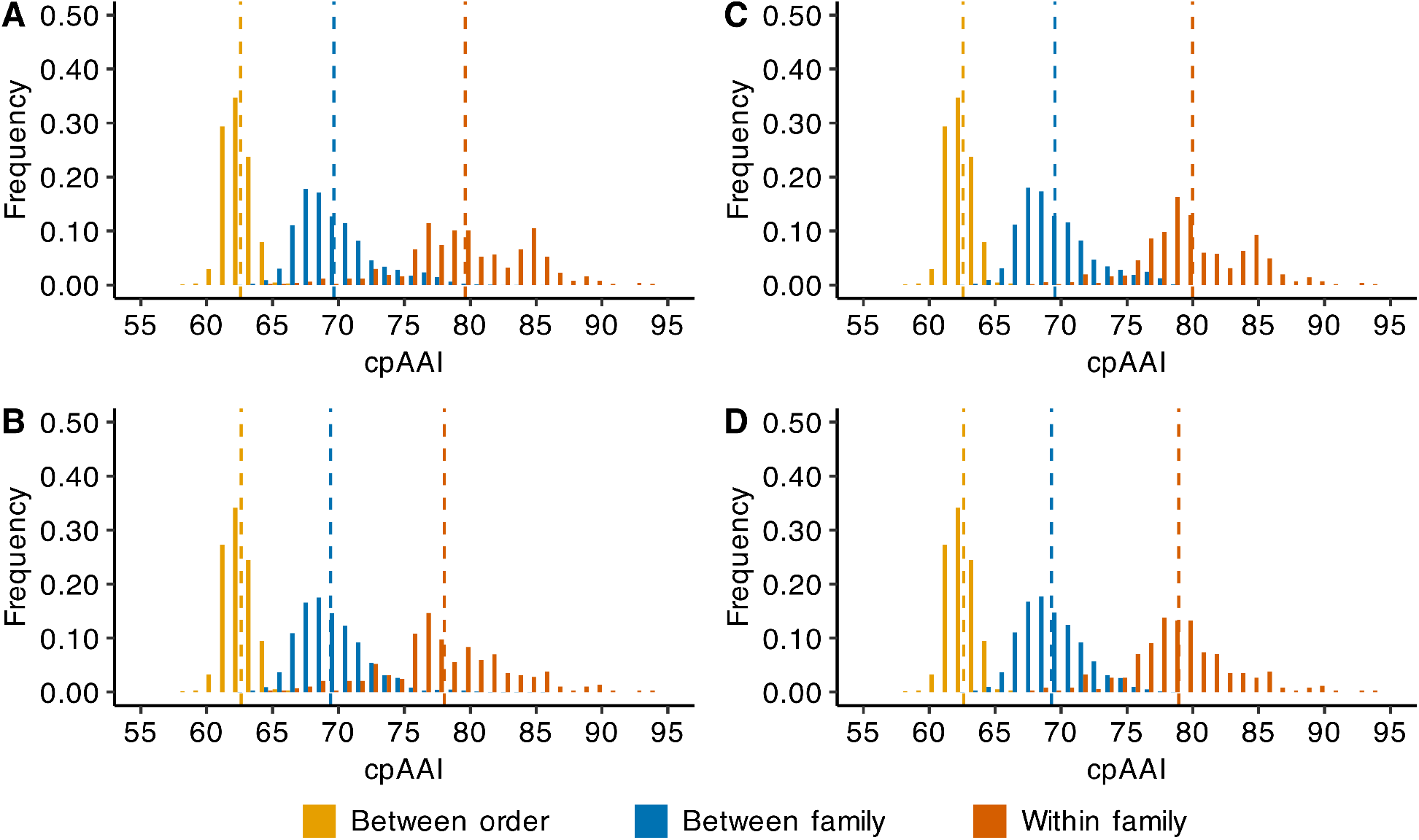
Distribution of core-proteome AAI (cpAAI) comparisons of the order *Hyphomicrobiales*. Pairwise cpAAI values were calculated based on 59 nonrecombinant loci from the core genome of 133 members of the order *Hyphomicrobiales* and five members of the order *Caulobacterales*. Results are summarized as histograms with a bin width of 1%. The cpAAI values calculated between two strains belonging to different orders (yellow), different families but same order (blue), or the same family (red) are summarized separately. Dashed vertical lines represent the mean value of each distribution. In all plots, cpAAI values where both strains belong to the order *Caulobacterales* were excluded. (A) The distribution of all pairwise cpAAI values with the classification (i.e., between order, between family, or within family) based on existing taxonomic assignments. (**B**) The distribution of all pairwise cpAAI values except for those including at least one strain from the family *Rhizobiaceae*, with the classification based on existing taxonomic assignments. (**C**) The distribution of all pairwise cpAAI values with the classification based on the proposed taxonomic assignments. (**D**) The distribution of all pairwise cpAAI values except for those including at least one strain from the family *Rhizobiaceae*, with the classification based on the proposed taxonomic assignments.

The within-family, between-family, and between-order distributions of cpAAI were clearly distinct, with mean values of 78.0%, 69.4%, and 62.6%, respectively, for the dataset lacking the family *Rhizobiaceae*. Notably, excluding the family *Rhizobiaceae*, 97.1% of the between-family comparisons fell below 75%, while 80.1% of the within-family comparisons were above 75% (**Figure 2B**). When considering the taxonomic changes proposed below, this increases to 98.3% of between-family comparisons being below 75% and 88.5% (93.8% when excluding *Bartonella bacilliformis* KC583^T^ that has a highly reduced genome) of within-family comparisons being above 75% (**Figure 2D**). These results, together with the shapes of the distributions, suggested to us that a cpAAI value of ∼75%, when calculated from core_138, represented a natural threshold for defining families within the order *Hyphomicrobiales*. Performing similar analyses using cpAAI values calculated from core_143 (**Figure S5**), or the wpAAI data (**Figure S6**), also allowed for the identification of potential thresholds (∼78% and ∼63%, respectively) for defining families within the order *Hyphomicrobiales*. However, these thresholds were less well defined compared to the threshold identified for core_138 and thus we focus our analyses primarily on the cpAAI data generated with core_138.

We attempted to supplement our genome-based analyses with 16S rRNA gene phylogenetic studies (**Figures S7 and S8**). However, the 16S rRNA gene phylogenies we built were non-concordant with the genome-based phylogenies and failed to accurately resolve multiple families, consistent with past studies [19]. We therefore only made use of the 16S rRNA gene phylogenies for taxa lacking whole genome sequencing data.

In the following sections, we propose several taxonomic revisions in the order *Hyphomicrobiales* based on overall genomic similarity and the need for families to be monophyletic. In general, we aimed to be conservative and limit the number of changes that we propose, and therefore have not proposed to split monophyletic families based solely on low cpAAI values. We do, however, propose to unify sister families where appropriate and only when clearly supported by the cpAAI data.

### Taxonomic implications for the families Rhizobiaceae, Ahrensiaceae, Bartonellaceae, Brucellaceae, Phyllobacteriaceae, and Notoacmeibacteraceae

The six families *Rhizobiaceae*, *Ahrensiaceae*, *Bartonellaceae*, *Brucellaceae*, *Phyllobacteriaceae*, and *Notoacmeibacteraceae* collectively represent over a third of the *Hyphomicrobiales* strains included in our analyses. Of note was the family *Phyllobacteriaceae*, which was not monophyletic in any of the phylogenies and whose genus type strain (*Phyllobacterium myrsinacearum* DSM 5892^T^) consistently formed its own lineage separate from all other members of the family (**Figures 1, S1, S2, and S4**). *Pseudohoeflea suaedae* YC6898^T^ was consistently nested within the family *Rhizobiaceae* in all phylogenies (**Figures 1, S1, S2, and S4**). We therefore propose that the genus *Pseudohoeflea* be reassigned to the family *Rhizobiaceae*. In addition, *Salaquimonas pukyongi* RR3-28^T^ consistently formed its own lineage (**Figures 1, S1, S2, and S4**), and when calculated from core_138, displayed pairwise cpAAI values <74% against all other strains (**Figure 1**). We therefore propose that the genus *Salaquimonas* be reassigned to the family *Salaquimonadaceae* fam. nov.. Moreover, *Oricola cellulosilytica* JCM 19534^T^, *Roseitalea porphyridii* MA7-20^T^, and *Oceaniradius stylonematis* StC1^T^ consistently grouped separately from the rest of the family *Phyllobacteriaceae* and instead formed a clade with *Ahrensia kielensis* DSM 5890^T^ (**Figures 1, S1, S2, and S4**), the genus type strain for the type genus of the family *Ahrensiaceae*. When calculated from core_138, all pairwise cpAAI values between the four strains were >78% (**Figure 1**). We therefore propose that the genera *Oricola*, *Roseitalea*, and *Oceaniradius* be reassigned to the family *Ahrensiaceae*.

The remaining 11 members of the family *Phyllobacteriaceae* included in this analysis were paraphyletic in all four phylogenies (**Figures 1, S1, S2, and S4**). The minimal monophyletic group containing all 11 *Phyllobacteriaceae* species also includes the families *Brucellaceae*, *Bartonellaceae*, and *Notoacmeibacteraceae*; the inclusion of the genus *Bartonella* within this monophyletic group was further supported by supplemental phylogenetic analyses (see **Text S2**; **Figures S9-S12**). As a result, it was necessary either to split the family *Phyllobacteriaceae* into three families or to unify the families *Phyllobacteriaceae*, *Brucellaceae*, *Bartonellaceae*, and *Notoacmeibacteraceae*. The cpAAI and wpAAI measurements favour the latter option; when calculated from core_138, all pairwise cpAAI values between members of these families were above 76.5%, except for comparisons involving *B. bacilliformis* KC583^T^, which has a highly reduced genome (cpAAI values involving this strain ranged from 69% to 74.5%), or *Notoacmeibacter marinus* XMTR2A4^T^, which displayed unexpectedly low cpAAI values against all strains (cpAAI values involving this strain ranged from 69% to 76%). Likewise, most cpAAI comparisons calculated with core_143 and wpAAI values were above the proposed thresholds of 78% and 63%, respectively. Based on these results, we propose that the families *Bartonellaceae*, *Brucellaceae*, *Notoacmeibacteraceae* and *Phyllobacteriaceae* be unified with the name *Bartonellaceae* as it takes priority. In addition, the strain *Limoniibacter endophyticus* KCTC 42097^T^, currently assigned to the family *Hyphomicrobiaceae*, was consistently nested within the emended family *Bartonellaceae* in all phylogenies. We therefore propose that the genus *Limoniibacter* be reassigned to the family *Bartonellaceae*.

In contrast to our initial expectations [4], the data suggested that the family *Rhizobiaceae* did not need to be split into two or more families, while our supplemental phylogenetic analyses (see **Text S2**; **Figures S9-S12**) supported the inclusion of the recently proposed genus “*Flavimaribacter*” in the family *Rhizobiaceae* [20]. On the other hand, when calculated from core_138, 82.5% of the cpAAI comparisons between members of the family *Rhizobiaceae* and the emended family *Bartonellaceae* were above our proposed threshold of 75% (**Figure 1**). Similarly, 71.2% of the wpAAI comparisons between members of the family *Rhizobiaceae* and the emended family *Bartonellaceae* were above our proposed threshold of 63% (**Figure S3**), although only 36.6% of the cpAAI comparisons were above our proposed threshold of 78% when calculated from core_143 (**Figure S1**). The possibility of uniting these two families therefore merits consideration. However, we do not favour this approach for two reasons. First, the between-family cpAAI values (calculated from core_138) tended to be lower than the within-family cpAAI values (**Figure 1**), with average values of 75.9% and 80.5%, respectively; qualitatively similar results are seen with cpAAI calculated from core_143 and the wpAAI data (**Figure S1 and S3**). In addition, in three of the four phylogenies, the families *Rhizobiaceae* and *Bartonellaceae* were not sister taxa; instead, these families only formed a monophyletic group when the emended family *Ahrensiaceae* was also included. The family *Ahrensiaceae* was better delineated from the family *Rhizobiaceae* based on cpAAI and wpAAI values (**Figures 1, S1, and S3**); the average cpAAI value (when calculated from core_138) for comparisons between species of the families *Ahrensiaceae* and *Rhizobiaceae* was 74.1%, with 84% of comparisons below the proposed threshold of ∼75% for family delineation. Likewise, in the cases of cpAAI calculated from core_143 and wpAAI, 99% and 86%, respectively, of the comparisons between species of the families *Ahrensiaceae* and *Rhizobiaceae* were below the proposed thresholds for family delineation. These results suggest it would not be appropriate to unify the families *Ahrensiaceae* and *Rhizobiaceae*. Considering the weight of the evidence, we have therefore chosen to leave the *Rhizobiaceae*, the emended *Ahrensiaceae*, and the emended *Bartonellaceae* as distinct families.

### Taxonomic implications: formation of new families

Several families in the order *Hyphomicrobiales* were paraphyletic in all four of our phylogenies. To fix the paraphyly and ensure all families are monophyletic, we propose the formation of three new families and the reassignment of three genera to other existing families (see the following section).

The genus type strain *Rhodoblastus acidophilus* DSM 137^T^ is currently assigned to the family *Roseiarcaceae*. However, *R. acidophilus* DSM 137^T^ did not form a monophyletic group with the genus type strain *Roseiarcus fermentans* DSM 24875^T^ that represents the type genus of the family *Roseiarcaceae*. Instead, *R. acidophilus* DSM 137^T^ formed its own lineage as a sister taxon to the family *Methylocystaceae* in all four phylogenies (**Figures 1, S1, S2, and S4**). When calculated from core_138, the pairwise cpAAI values between *R. acidophilus* DSM 137^T^ and the two included genus type strains of the family *Methylocystaceae* were 74.8% and 74.5% (**Figure 1**), which are below our proposed threshold of 75% for family deliminatation. We therefore propose to reassign the genus *Rhodoblastus* to the family *Rhodoblastaceae* fam. nov..

The genus *Methylopila* was not assigned to a family when it was proposed [21], but was subsequently assigned to the family *Methylocystaceae* [22]. However, the genus type strain *Methylopila capsulata* DSM 6130^T^ did not cluster with the other members of the family *Methylocystaceae*, and instead formed its own distinct lineage in the two phylogenies based on core_138 or perc95_138 (**Figures 1 and S3**). When calculated from core_138, all comparisons involving *M. capsulata* DSM 6130^T^ gave cpAAI values less than 74% (**Figure 1**). In the phylogenies based on core_143 or perc95_143, which include five additional genus type strains, *M. capsulata* DSM 6130^T^ formed a clade with *Chenggangzhangella methanolivorans* CHL1^T^ also currently assigned to the family *Methylocystaceae* (**Figures S1 and S2**). When calculated from core_143, these two strains had a pairwise cpAAI value of 86.5% (**Figure S1**), which is above the proposed threshold of 78% for family delineation. The 16S rRNA gene phylogenies (**Figures S7 and S8**) included two additional genus type strains, *Hansschlegelia plantiphila* VKM B-2347^T^ and *Albibacter methylovorans* DM10^T^, of genera currently assigned to the family *Methylocystaceae* but that lack publicly available whole genome sequence. In both phylogenies, *H. plantiphila* VKM B-2347^T^ and *A. methylovorans* DM10^T^ formed a monophyletic clade with *M. capsulata* DSM 6130^T^ and *C. methanolivorans* CHL1^T^, rather than with *Methylocystis parvus* OBBP^T^, the genus type strain of the type genus of the family *Methylocystaceae*. Based on these results, we propose to reassign the genera *Methylopila*, *Chenggangzhangella*, *Hansschlegelia*, and *Albibacter* to the family *Methylopilaceae* fam. nov..

The genus type strain *Rhodoligotrophos appendicifer* 120-1^T^ is currently assigned to the family *Parvibaculaceae*. However, instead of clustering with the other members of the family *Parvibaculaceae*, it formed its own lineage as a sister taxon to the genus type strain *Aestuariivirga litoralis* KCTC 52945^T^ of the family *Aestuariivirgaceae* in all four phylogenies (**Figures 1, S1, S2, and S4**). When calculated from core_138, the pairwise cpAAI value between *R. appendicifer* 120-1^T^ and *A. litoralis* KCTC 52945^T^ was 71.2% (**Figure 1**), which is below our proposed threshold of 75% for family delineation. We therefore propose to reassign the genus *Rhodoligotrophos* to the family *Rhodoligotrophaceae* fam. nov..

### Taxonomic implications: reassignment of genera

The genus type strain *Faunimonas pinastri* A52C2^T^ is currently assigned to the family *Pleomorphomonadaceae* but did not cluster with other members of this family. Instead, in all four phylogenies, *F. pinastri* A52C2^T^ formed a clade with the genus type strain *Afifella marina* DSM2698^T^ that represents the only genus of the family *Afifellaceae* (**Figures 1, S1, S2, and S4**). When calculated from core_138, the cpAAI value between *F. pinastri* A52C2^T^ and *A. marina* DSM2698^T^ was 75.6% (**Figure 1**), which is above our proposed threshold of 75% for family delimitation. We therefore propose to reassign the genus *Faunimonas* to the family *Afifellaceae*.

The genus *Pseudochelatococcus* was not assigned to a family when it was proposed [23], but was subsequently associated with the family *Beijerinckiaceae* [24]. However, the genus type strain *Pseudochelatococcus lubricantis* DSM 103870^T^ did not cluster with the other members of the family *Beijerinckiaceae*, and instead formed a clade with *Chelatococcus asaccharovorans* DSM 6462^T^ and *Camelimonas lactis* DSM22958^T^ from the family *Chelatococcaceae* in all four phylogenies (**Figures 1, S1, S2, and S4**). Moreover, the family *Chelatococcaceae* was not monophyletic without the inclusion of *P. lubricantis* DSM 103870^T^ (**Figures 1, S1, S2, and S4**). When calculated from core_138, all pairwise cpAAI values in this clade were >75% (**Figure 1**). We therefore propose to assign the genus *Pseudochelatococcus* to the family *Chelatococcaceae*.

The genus type strain *Prosthecomicrobium pneumaticum* DSM 16268^T^ was previously assigned to the family *Hyphomicrobiaceae* [25], although it was recently proposed that the genus *Prosthecomicrobium* be reassigned to the family *Kaistiaceae* [26]. In all four phylogenies, *P. pneumaticum* DSM 16268^T^ formed a clade with *Kaistia adipata* DSM 17808^T^ and *Bauldia litoralis* ATCC 35022^T^ of the family *Kaistiaceae* rather than with species of the family *Hyphomicrobiaceae* (**Figures 1, S1, S2, and S4**). Moreover, the family *Kaistiaceae* was not monophyletic without the inclusion of *P. pneumaticum* DSM 16268^T^ (**Figures 1, S1, S2, and S4**). When calculated from core_138, all pairwise cpAAI values in this clade were >75% (**Figure 1**). Our data are therefore consistent with the proposal to reassign the genus *Prosthecomicrobium* to the family *Kaistiaceae*.

### Taxonomic implications: unification of families

The families *Segnochrobactraceae* and *Pseudoxanthobacteraceae* each contain a single validly published genus. The genus type strains of these two genera formed a monophyletic group in all four phylogenies (**Figures 1, S1, S2, and S4**). When comparing the genomes of these two strains, the pairwise cpAAI value was 82.9% when calculated from core_138 (**Figure 1**) and 85.9% when calculated from core_143 (**Figure S1**), while the wpAAI value was 70.0% (**Figure S3**). Given that these values are all well above the proposed thresholds for family delimitation, we propose that the families *Segnochrobactraceae* and *Pseudoxanthobacteraceae* be unified with the name *Segnochrobactraceae* as it takes priority.

The families *Lichenihabitantaceae* and *Lichenibacteriaceae* each contain a single validly published genus, based on strains that were originally isolated from lichens in the Antarctic and the subarctic zone of the northern hemisphere, respectively [27, 28]. The genus type strains of these two genera formed a monophyletic group in all four phylogenies (**Figures 1, S1, S2, and S4**). When comparing the genomes of these two strains, the pairwise cpAAI value was 77.9% when calculated from core_138 (**Figure 1**) and 78.9% when calculated from core_143 (**Figure S1**), while the wpAAI value was 67% (**Figure S3**). Given that these values are all above the proposed thresholds for family delimitation, we propose that the families *Lichenihabitantaceae* and *Lichenibacteriaceae* be unified with the name *Lichenihabitantaceae* as it takes priority.

The family *Breoghaniaceae* contains a single genus while the family *Stappiaceae* contains five genera. The six genus type strains from these two families formed a monophyletic group in all four phylogenies (**Figures 1, S1, S2, and S4**). When calculated from core_138, the pairwise cpAAI values between members of the family *Stappiaceae* ranged between 77.0% and 81.6%, which overlapped the cpAAI range of 76.2% to 78.9% for comparisons between members of the family *Stappiaceae* and *Breoghania corrubedonensis* DSM 23382^T^ (**Figure 1**). Similar results are seen when using cpAAI calculated from core_143 and wpAAI values (**Figures S1 and S3**). Considering these results, we propose that the families *Stappiaceae* and *Breoghaniaceae* be unified with the name *Stappiaceae*. Both family names were published in the same article [2] but *Stappiaceae* is our preferred choice because *Breoghaniaceae* includes only a single genus.

### Other taxonomic implications

In this study, we refrained from splitting monophyletic families based solely on cpAAI data. Nevertheless, multiple families merit further study as candidates for splitting, namely the families *Amorphaceae*, *Devosiaceae*, *Hyphomicrobiaceae*, and *Parvibaculaceae* (**Figure 1**). Additionally, a couple of clades were not well-resolved based on the cpAAI data and the proposed thresholds, and thus also merit further study. The family *Rhodobiaceae* and the emended family *Stappiaceae* are sister taxa that are not well separated in the cpAAI data calculated from core_138 (**Figure 1**), with four of the six between-family comparisons above 75%, although better separation is observed in the cpAAI data calculated from core_143 and the wpAAI data (**Figure S1 and S3**). Likewise, the families *Methylobacteriaceae* and *Salinarimonadaceae* were not well separated by the cpAAI or wpAAI data; six of the eight cpAAI values calculated from core_138 were above 75% with the other two above 74.7% (**Figure 1**), four of the eight cpAAI values calculated from core_143 were above 78% with the other four above 77.5% (**Figure S1**), and all wpAAI values were above the proposed threshold of 63% (**Figure S3**). As there was not overwhelming support for unification of the families *Rhodobiaceae* and *Stappiaceae*, or the families *Methylobacteriaceae* and *Salinarimonadaceae*, we chose to leave all four as distinct families. However, these taxa merit further consideration in future work.

Although it is not a genus type strain, we included the species type strain “*Pararhizobium mangrovi*” BGMRC 6574^T^ in our dataset as preliminary investigations of the family *Rhizobiaceae* suggested it belonged to a novel genus and potentially a novel family. Our full analysis did not support the reassignment of “*P. mangrovi*” BGMRC 6574^T^ to a new family but it did support the recent proposal to reclassify this strain as “*Allopararhizobium mangrovi*” BGMRC 6574^T^ once validly published [29].

### Emended description of *Hyphomicrobiales* Douglas et al. 1957 (Approved Lists 1980) emend. Hördt et al. 2020

The description is as given before [1, 2], with the following modification. The order currently comprises the families *Aestuariivirgaceae*, *Afifellaceae*, *Ahrensiaceae*, *Alsobacteraceae*, *Amorphaceae*, *Ancalomicrobiaceae*, *Aurantimonadaceae*, *Bartonellaceae*, *Beijerinckiaceae*, *Blastochloridaceae*, *Boseaceae*, *Chelatococcaceae*, *Cohaesibacteraceae*, *Devosiaceae*, *Hyphomicrobiaceae*, *Kaistiaceae*, *Lichenihabitantaceae*, *Methylobacteriaceae*, *Methylocystaceae*, *Methylopilaceae* fam. nov., *Nitrobacteraceae*, *Parvibaculaceae*, *Phreatobacteraceae*, *Pleomorphomonadaceae*, *Rhabdaerophilaceae*, *Rhizobiaceae*, *Rhodobiaceae*, *Rhodoblastaceae* fam. nov., *Rhodoligotrophaceae* fam. nov., *Roseiarcaceae*, *Salaquimonadaceae* fam. nov., *Salinarimonadaceae*, *Segnochrobactraceae*, *Stappiaceae*, *Tepidamorphaceae*, and *Xanthobacteraceae*. The type genus is *Hyphomicrobium*. The families *Breoghaniaceae*, *Brucellaceae*, *Lichenibacteriaceae*, *Notoacmeibacteraceae*, *Phyllobacteriaceae*, and *Pseudoxantobacteraceae* were removed from the order and their genera transferred to other families within the order.

### Description of *Methylopilaceae* fam. nov

Me.thyl.o.pil.a’ce.ae (N.L. fem. n. *Methylopila*, type genus of the family; -*aceae*, ending to denote a family; N.L. fem. pl. n. *Methylopilaceae*, the *Methylopila* family).

Cells are aerobic, Gram-negative, rod-shaped or cocci, and motile or non-motile. The G+C content as calculated from genome sequences is 68.2-69.7%, while the range provided in the literature is 64.4-70.4 mol% [30, 31]. The family currently comprises the genera *Albibacter*, *Chenggangzhangella*, *Hansschlegelia*, and *Methylopila* (the type genus). This family can be differentiated from other families based on phylogenetic analyses of core proteome and 16S rRNA gene sequences, as well as OGRI calculations (cpAAI and wpAAI).

### Description of *Rhodoblastaceae* fam. nov

Rho.do.blast.a’ceae (N.L. masc. n. *Rhodoblastus*, type genus of the family; -*aceae*, ending to denote a family; N.L. fem. pl. n. *Rhodoblastaceae*, the *Rhodoblastus* family).

The description is as given for *Rhodoblastus* [32], which is the type and currently the sole genus of the family. This family can be differentiated from other families based on phylogenetic analyses of core proteome sequences and OGRI calculations (cpAAI and wpAAI).

### Description of *Rhodoligotrophaceae* fam. nov

Rho.do.li.go.tro’ph.a’ce.ae (N.L. masc. n. *Rhodoligotrophos*, type genus of the family; -*aceae*, ending to denote a family; N.L. fem. pl. n. *Rhodoligotrophaceae*, the *Rhodoligotrophos* family).

The description is as given for *Rhodoligotrophos* [33], which is the type and currently the sole genus of the family. This family can be differentiated from other families based on phylogenetic analyses of core proteome sequences and OGRI calculations (cpAAI and wpAAI).

### Description of *Salaquimonadaceae* fam. nov

Sal.a.qui.mo’nad.a’ce.ae (N.L. fem. n. *Salaquimonas*, type genus of the family; -*aceae*, ending to denote a family; N.L. fem. pl. n. *Salaquimonadaceae*, the *Salaquimonas* family).

The description is as given for *Salaquimonas* [34], which is the type and currently the sole genus of the family. This family can be differentiated from other families based on phylogenetic analyses of core proteome sequences and OGRI calculations (cpAAI and wpAAI).

### Emended description of *Afifellaceae* Hördt et al. 2020

The description is as given before [2], with the following modification. The family currently comprises the genera *Afifella* (the type genus), and *Faunimonas*.

### Emended description of *Ahrensiaceae* Hördt et al. 2020

The description is as given before [2], with the following modification. The family currently comprises the genera *Ahrensia* (the type genus), *Pseudahrensia*, *Oricola*, *Roseitalea*, and *Oceaniradius*.

### Emended description of *Bartonellaceae* Gieszczykiewicz 1939 (Approved Lists 1980) emend. Brenner et al. 1993

The description is based on descriptions provided previously [2, 35–39]. Cells are Gram-negative, aerobic, with variable morphology (rod, ovoid, coccoid, coccobacilli, or ring or disk shaped). Cells can be motile by means of flagella, or non-motile. Generally catalase and oxidase positive. Predominantly aerobic or facultatively anaerobic heterotrophs, with some species able to utilize carbohydrates while others cannot. Generally non-spore-forming. The optimum growth temperature ranges between 20°C and 37°C. The predominant respiratory quinone is generally Q-10. Some species are human pathogens. The G+C content as calculated from genome sequences of genus type strains is 38.2-65.1%

The family currently comprises the genera *Aminobacter*, *Aquamicrobium*, *Aquibium*, *Bartonella* (the type genus), *Brucella*, *Chelativorans*, *Corticibacterium*, *Daeguia*, *Falsochrobactrum*, *Limoniibacter*, *Mesorhizobium*, *Nitratireductor*, *Notoacmeibacter*, *Ochrobactrum*, *Paenochrobactrum*, *Phyllobacterium*, *Pseudaminobacter*, *Pseudochrobactrum*, *Tianweitania*, and *Zhengella*.

### Emended description of *Beijerinckiaceae* Grarity et al. 2006 emend. Hördt et al. 2020

The description is as given before [2]. The family currently comprises the genera *Beijerinckia* (the type genus), *Methylocapsa*, *Methylocella*, *Methyloferula*, *Methylorosula*, and *Methylovirgula*. The genus *Pseudochelatococcus* was removed from the family and transferred to the family *Chelatococcaceae*.

### Emended description of *Chelatococcaceae* Dedysh et al. 2016

The description is as given before [40], with the following modification. The family currently comprises the genera *Camelimonas*, *Chelatococcus* (the type genus), and *Pseudochelatococcus*.

### Emended description of *Hyphomicrobiaceae* Babudieri 1950 (Approved Lists 1980) emend. Hördt et al. 2020

The description is as given before [2] with the following modifications. The family currently comprises the genera *Dichotomicrobium*, *Filomicrobium*, *Hyphomicrobium* (the type genus), *Methyloceanibacter*, *Methyloligella*, *Pedomicrobium*, *Rhodomicrobium*, and *Seliberia*. The genus *Limoniibacter* was removed from the family and transferred to the emended family *Bartonellaceae*. The family also does not encompass the genus *Caenibius* (*Erythrobacteraceae, Sphingomonadales*), which appears to have been mistakenly included in the description of the family *Hyphomicrobiaceae* by Hördt et al. (2020) [2].

### Emended description of *Lichenihabitantaceae* Noh et al. 2019

The description is as given before [27] with the following modifications. Cells are aerobic or facultative anaerobic. The family currently comprises the genera *Lichenibacterium* and *Lichenihabitans* (the type genus).

### Emended description of *Methylocystaceae* Bowman 2006 emend. Hördt et al. 2020

The description is as given before [2]. The family currently comprises the genera *Methylocystis* (the type genus) and *Methylosinus*. The genera *Albibacter*, *Chenggangzhangella*, *Hansschlegelia*, and *Methylopila* were removed from the family and transferred to *Methylopilaceae* fam. nov..

### Emended description of *Parvibaculaceae* Hördt et al. 2020

The description is as given before [2], with the following modification. The family currently comprises the genera *Parvibaculum* (the type genus) and *Tepidicaulis*. The genera *Anderseniella* and *Pyruvatibacter* are tentatively assigned to this family as well. The genus *Rhodoligotrophos* was removed from the family and transferred to *Rhodoligotrophaceae* fam. nov..

### Emended description of *Pleomorphomonadaceae* Hördt et al. 2020

The description is as given before [2]. The family currently comprises the genera *Chthonobacter*, *Hartmannibacter*, *Methylobrevis*, *Mongoliimonas*, *Oharaeibacter*, and *Pleomorphomonas* (the type genus). The genus *Faunimonas* assigned to this family by Proença et al. (2022) [41] was removed from the family and transferred to the family *Afifellaceae*.

### Emended description of *Rhizobiaceae* Conn 1938 (Approved Lists 1980) emend. Hördt et al. 2020

The description is as given before [2, 42], with the following modification. The family currently comprises the genera *Agrobacterium*, *Allorhizobium*, *Ciceribacter*, *Endobacterium*, *Ensifer*, *Ferranicluibacter*, *Gellertiella*, *Georhizobium*, *Hoeflea*, *Lentilitoribacter*, *Liberibacter*, *Martelella*, *Mycoplana*, *Neorhizobium*, *Onobrychidicola*, *Pararhizobium*, *Peteryoungia*, *Pseudohoeflea*, *Pseudorhizobium*, *Rhizobium* (the type genus), *Shinella*, *Sinorhizobium*, and *Xaviernesmea*.

### Emended description of *Roseiarcaceae* Kulichevskaya et al. 2014 emend. Hördt et al. 2020

The description is as given before [2, 43], with the following modification. The family comprises the genus *Roseiarcus*, which is the type and currently the sole genus of the family. The genus *Rhodoblastus* was removed from the family and transferred to *Rhodoblastaceae* fam. nov..

### Emended description of *Segnochrobactraceae* Akter et al. 2020

The description is as given before [44], with the following modification. The family currently comprises the genera *Pseudoxanthobacter* and *Segnochrobactrum* (the type genus).

### Emended description of *Stappiaceae* Hördt et al. 2020

The description is as given before [2], with the following modification. The family currently comprises the genera *Breoghania*, *Hongsoonwoonella*, *Pannonibacter*, *Pseudovibrio*, *Roseibium*, and *Stappia* (the type genus).

## ACKNOWLEDGEMENTS

This work was supported by a Natural Sciences and Engineering Research Council of Canada (NSERC) grant to G.C.D.

