## Supplementary material for "Refining the taxonomy of the order *Hyphomicrobiales* (*Rhizobiales*) based on whole genome comparisons of over 130 genus type strains": File S1

**This PDF file includes:**

Text S1 to S2 (pages 2-3)
Figures S1 to S12 (pages 4-21)
Legends for Datasets S1 to S17 (page 22)

**Other supplementary materials for this manuscript include the following:**

Datasets S1 to S17

**Text S1**

Preliminary investigation of the perc95_143 gene set (i.e., the 256 marker genes identified in at least 95% of the 143 strains included in our dataset) identified five strains whose publicly available genome lacked >10% of these genes. The low perc95_143 prevalence in *Liberibacter crescens* BT-1$^{T}$ is likely a consequence of its highly reduced genome, although we note that *Bartonella bacilliformis* KC583$^{T}$ has a similar genome size but contained orthologs of all perc95_143 genes. The low perc95_143 prevalence in *Methylobrevis pamukkalensis* VKM b-2849$^{T}$, *Chenggangzhangella methanolivorans* CHL1$^{T}$, and *Nitratireductor aquibiodomus* JCM 21793$^{T}$ is likely a consequence of the high percentage of pseudogenes in these genomes (16% of genes are annotated as pseudogenes). While this high rate of pseudogenes may be accurate, we cannot rule out that it was caused by errors in the genome assemblies. The reason for the low perc95_143 prevalence in *Methyloligella halotolerans* VKM B-2706$^{T}$ was less obvious. However, it may reflect a combination of the below average protein-coding gene count (2,872 vs an average of 4,294), the above average pseudogene rate (7.3% versus 2.5%), and the genome assembly being in contigs rather than fully assembled. Eliminating these five strains from our dataset more than tripled the number of non-recombining core genes from 19 to 59. As our dataset included at least one additional representative from each of the families containing these five strains, we repeated the phylogenetic and OGRI analyses without these strains to obtain more robust results.

**Text S2**

As members of the genus *Bartonella* have highly reduced genomes, we chose to perform a secondary analysis to be more confident that the genus *Bartonella* falls within the monophyletic group that includes the families *Phyllobacteriaceae*, *Notoacmeibacteraceae*, and *Brucellaceae*. To this end, we repeated our phylogenetic analyses using a taxonomically restricted subset of the *Hyphomicrobiales* dataset, including the families *Phyllobacteriaceae*, *Notoacmeibacteraceae*, *Brucellaceae*, *Bartonellaceae*, *Ahrensiaceae*, *Rhizobiaceae*, and *Aurantimonadaceae*. In addition, we supplemented our dataset with four additional *Bartonella* species type strains (*Bartonella apis* PEB0122$^{T}$, *Bartonella birtlesii* IBS 325$^{T}$, *Bartonella henselae* ATCC 49882$^{T}$, and *Bartonella krasnovii* OE 1-1$^{T}$) that represent the full taxonomic breadth of the genus [1], as well as "*Flavimaribacter sediminis*" WL0058$^{T}$ proposed to belong to the family *Rhizobiaceae*. This yielded a final dataset of 58 strains (Dataset S3).

GET_HOMOLOGUES and GET_PHYLOMARKERS were used to identify non-recombining single-copy marker genes present in all 58 genomes. This led to the identification of 120 core genes (termed core_58). In addition, GET_HOMOLOGUES was used to identify 454 single-copy marker genes present in at least 95% of the strains (termed perc95_58). Repeating these analyses with a reduced set of 56 genomes lacking *L. crescens* BT-1$^{T}$ and *N. aquibiodomus* JCM 21793$^{T}$ (see Text S1) led to the identification of 178 core genes present in all strains (termed core_56), and 497 genes present in at least 95% of the genomes (termed perc95_56). As described in the Methods section of the main manuscript for the *Hyphomicrobiales* dataset, the proteins encoded by the four gene sets were used to construct maximum likelihood phylogenies using IQ-TREE with the best scoring model (core_58: LG+F+R6; perc95_58: LG+F+R8; core_56: LG+F+R5; perc95_56: LG+F+R9).

The five species type strains of the genus *Bartonella* formed a monophyletic group in all four of the resulting phylogenies, as expected (Figures S9-S12). Importantly, all four phylogenies placed the genus *Bartonella* within the monophyletic group that also included the families *Phyllobacteriaceae*, *Notoacmeibacteraceae*, and *Brucellaceae* (Figures S9-S12) although the exact position differed compared to the position of *B. bacilliformis* KC583$^{T}$ in the full *Hyphomicrobiales* phylogenies (Figures 1, S1, S2, and S3). Collectively, these results are consistent with the proposal to unify the families *Bartonellaceae*, *Brucellaceae*, *Notoacmeibacteraceae*, and *Phyllobacteriaceae* under the name *Bartonellaceae*. In addition, the four supplemental phylogenies supported the placement of the recently proposed genus "*Flavimaribacter*" within the family *Rhizobiaceae* (Figures S9-S12).

1. **Gutiérrez R, Shalit T, Markus B, Yuan C, Nachum-Biala Y, *et al.*** *Bartonella kosoyi* sp. nov. and *Bartonella krasnovii* sp. nov., two novel species closely related to the zoonotic *Bartonella elizabethae*, isolated from black rats and wild desert rodent-fleas. *Int J of Syst Evol Microbiol* 2020;70:1656–1665.

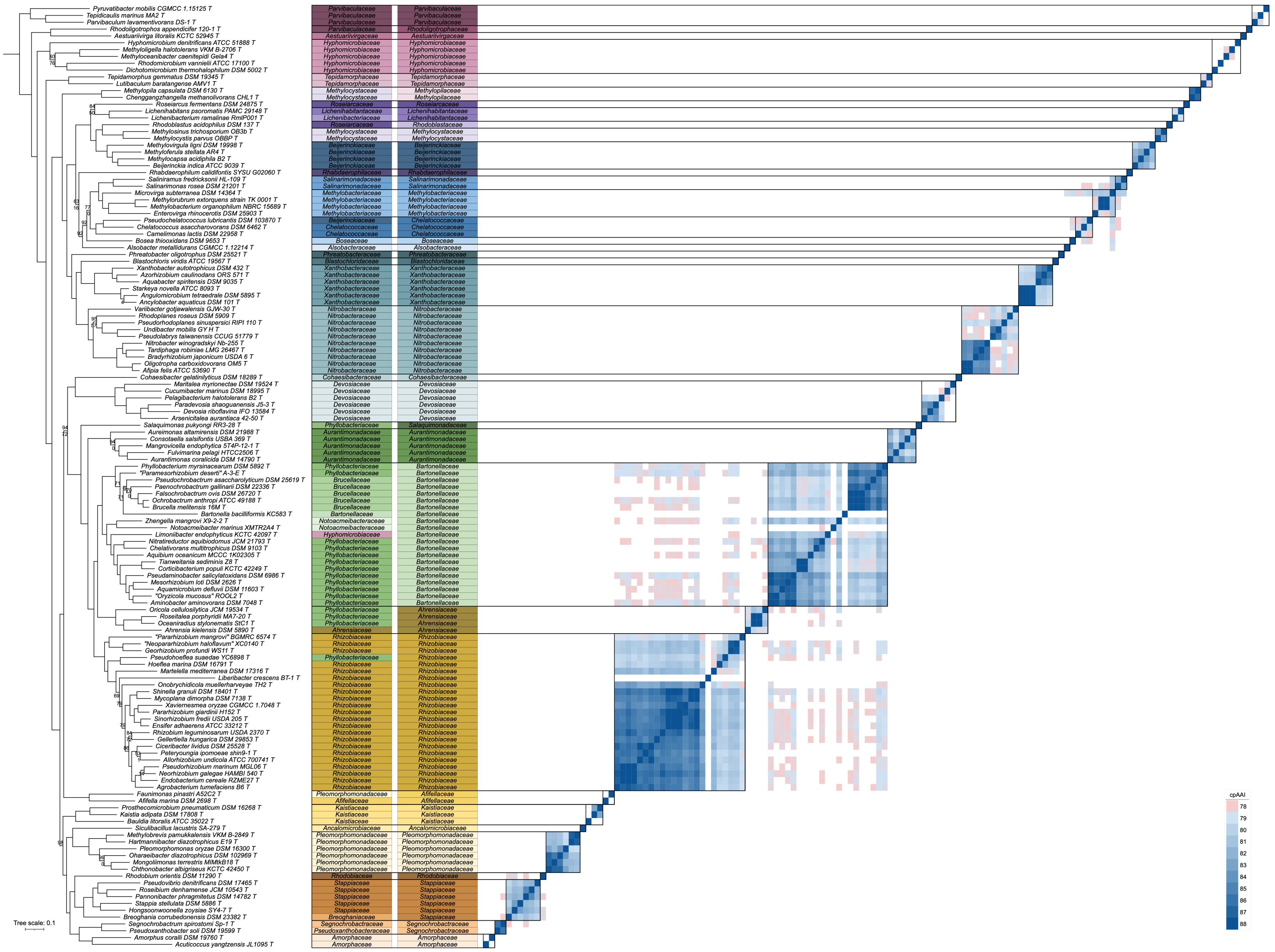
cpAAI
Tree scale: 0.1

**Figure S1. Phylogenetic and core-proteome AAI (cpAA) analyses of the order *Hyphomicrobiales*.** On the left, a maximum likelihood phylogeny of 138 *Hyphomicrobiales* type strains is shown, built using the concatenated protein alignments encoded by the perc95_143 gene set (256 genes present in at least 95% of the strains). The phylogeny was rooted using five *Caulobacterales* type strains as the outgroup. The numbers on the nodes indicate the ultra-fast jackknife values using a 40% resampling rate (top numbers) and the SH-aLRT support values (bottom numbers), both calculated from 1000 replicates. Only values below 100 are shown. The scale bar represents the average number of amino acid substitutions per site. To the right of the phylogeny is the current family assignments of each of the 138 *Hyphomicrobiales* type strains, followed to the right by the proposed family assignments of each strain. On the righthand side, a matrix is provided showing the cpAAI values between each pair of strains calculated using the proteins encoded by the core_143 gene set (19 genes present in 100% of the strains). Values less than 78% are in white while all values greater than 88% are the same shade of blue. Black boxes indicate the proposed families.

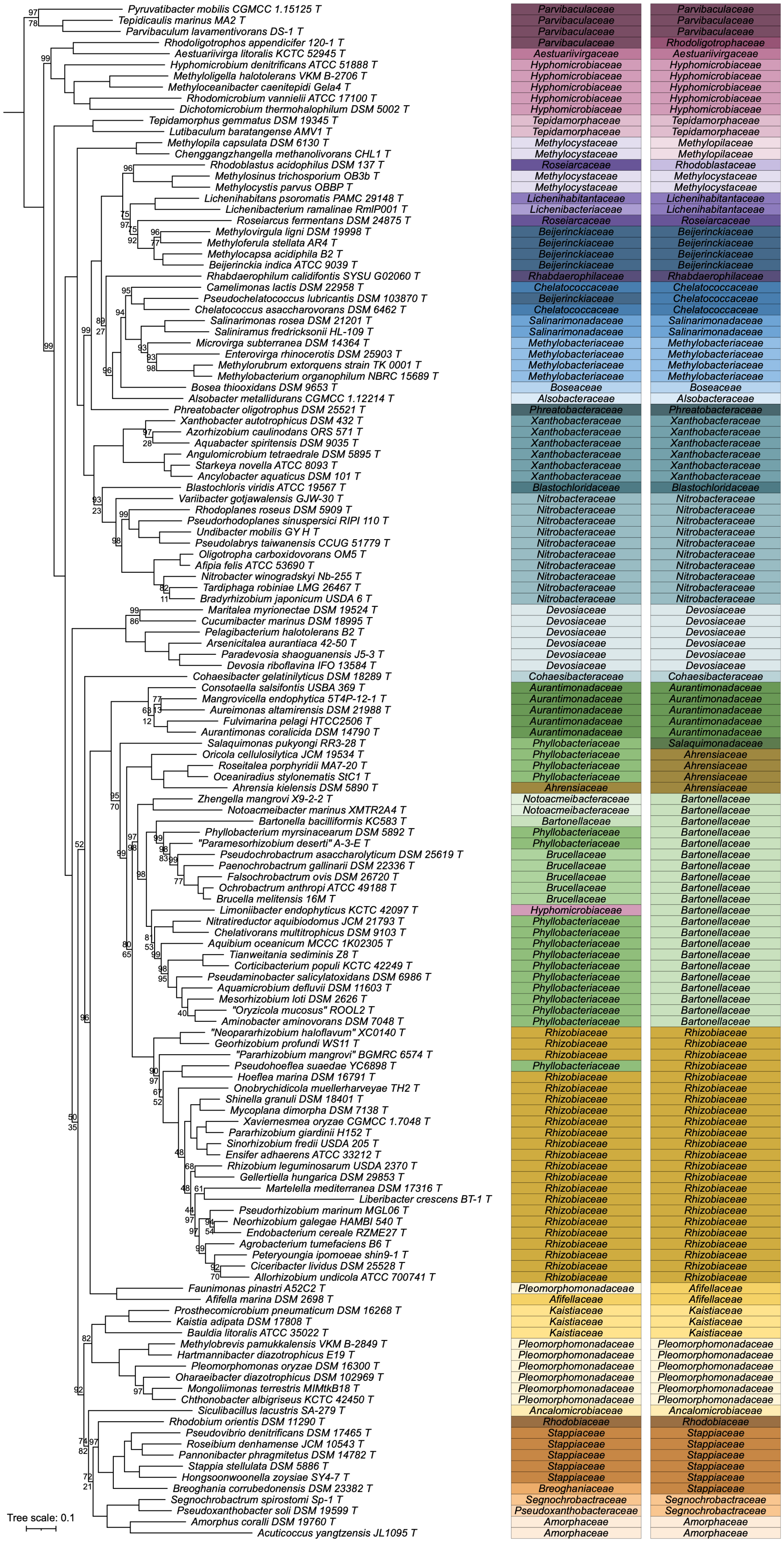
Pyruvatibacter mobilis CGMCC 1.15125 T Parvibaculaceae Parvibaculaceae
Tepidicaulis marinus MA2 T Parvibaculaceae Parvibaculaceae
Parvibaculum lavamentivorans DS-1 T Parvibaculaceae Parvibaculaceae
Rhodoligotrophos appendicifer 120-1 T Parvibaculaceae Rhodoligotrophaceae
Aestuariivirga litoralis KCTC 52945 T Aestuariivirgaceae Aestuariivirgaceae
Hyphomicrobium denitrificans ATCC 51888 T Hyphomicrobiaceae Hyphomicrobiaceae
Methyloligella halotolerans VKM B-2706 T Hyphomicrobiaceae Hyphomicrobiaceae
Methyloceanibacter caenitepidi Gela4 T Hyphomicrobiaceae Hyphomicrobiaceae
Rhodomicrobium vannielii ATCC 17100 T Hyphomicrobiaceae Hyphomicrobiaceae
Dichotomicrobium thermohalophilum DSM 5002 T Hyphomicrobiaceae Hyphomicrobiaceae
Tepidamorphus gemmatus DSM 19345 T Tepidamorphaceae Tepidamorphaceae
Lutibaculum baratangense AMV1 T Tepidamorphaceae Tepidamorphaceae
Methylopila capsulata DSM 6130 T Methylocystaceae Methylopilaceae
Chenggangzhangella methanolivorans CHL1 T Methylocystaceae Methylopilaceae
Rhodoblastus acidophilus DSM 137 T Roseiarcaceae Rhodoblastaceae
Methylosinus trichosporium OB3b T Methylocystaceae Methylocystaceae
Methylocystis parvus OBBP T Methylocystaceae Methylocystaceae
Lichenihabitans psoromatis PAMC 29148 T Lichenihabitantaceae Lichenihabitantaceae
Lichenibacterium ramalinae RmlP001 T Lichenibacteriaceae Lichenibacteriaceae
Roseiarcus fermentans DSM 24875 T Roseiarcaceae Roseiarcaceae
Methylovirgula ligni DSM 19998 T Beijerinckiaceae Beijerinckiaceae
Methyloferula stellata AR4 T Beijerinckiaceae Beijerinckiaceae
Methylocapsa acidiphila B2 T Beijerinckiaceae Beijerinckiaceae
Beijerinckia indica ATCC 9039 T Beijerinckiaceae Beijerinckiaceae
Rhabdaerophilum calidifontis SYSU G02060 T Rhabdaerophilaceae Rhabdaerophilaceae
Camelimonas lactis DSM 22958 T Chelatococcaceae Chelatococcaceae
Pseudochelatococcus lubricantis DSM 103870 T Beijerinckiaceae Chelatococcaceae
Chelatococcus asaccharovorans DSM 6462 T Chelatococcaceae Chelatococcaceae
Salinarimonas rosea DSM 21201 T Salinarimonadaceae Salinarimonadaceae
Saliniramus fredricksonii HL-109 T Salinarimonadaceae Salinarimonadaceae
Microvirga subterranea DSM 14364 T Methylobacteriaceae Methylobacteriaceae
Enterovirga rhinocerotis DSM 25903 T Methylobacteriaceae Methylobacteriaceae
Methylorubrum extorquens strain TK 0001 T Methylobacteriaceae Methylobacteriaceae
Methylobacterium organophilum NBRC 15689 T Methylobacteriaceae Methylobacteriaceae
Bosea thiooxidans DSM 9653 T Boseaceae Boseaceae
Alsobacter metallidurans CGMCC 1.12214 T Alsobacteraceae Alsobacteraceae
Phreatobacter oligotrophus DSM 25521 T Phreatobacteraceae Phreatobacteraceae
Xanthobacter autotrophicus DSM 432 T Xanthobacteraceae Xanthobacteraceae
Azorhizobium caulinodans ORS 571 T Xanthobacteraceae Xanthobacteraceae
Aquabacter spiritensis DSM 9035 T Xanthobacteraceae Xanthobacteraceae
Angulomicrobium tetraedrale DSM 5895 T Xanthobacteraceae Xanthobacteraceae
Starkeya novella ATCC 8093 T Xanthobacteraceae Xanthobacteraceae
Ancylobacter aquaticus DSM 101 T Xanthobacteraceae Xanthobacteraceae
Blastochloris viridis ATCC 19567 T Blastochloridaceae Blastochloridaceae
Variibacter gotjawalensis GJW-30 T Nitrobacteraceae Nitrobacteraceae
Rhodoplanes roseus DSM 5909 T Nitrobacteraceae Nitrobacteraceae
Pseudorhodoplanes sinuspersici RIPI 110 T Nitrobacteraceae Nitrobacteraceae
Undibacter mobilis GY H T Nitrobacteraceae Nitrobacteraceae
Pseudolabrys taiwanensis CCUG 51779 T Nitrobacteraceae Nitrobacteraceae
Oligotropha carboxidovorans OM5 T Nitrobacteraceae Nitrobacteraceae
Afipia felis ATCC 53690 T Nitrobacteraceae Nitrobacteraceae
Nitrobacter winogradskyi Nb-255 T Nitrobacteraceae Nitrobacteraceae
Tardiphaga robiniae LMG 26467 T Nitrobacteraceae Nitrobacteraceae
Bradyrhizobium japonicum USDA 6 T Nitrobacteraceae Nitrobacteraceae
Maritalea myrionectae DSM 19524 T Devosiaceae Devosiaceae
Cucumibacter marinus DSM 18995 T Devosiaceae Devosiaceae
Pelagibacterium halotolerans B2 T Devosiaceae Devosiaceae
Arsenicitalea aurantiaca 42-50 T Devosiaceae Devosiaceae
Paradevosia shaoguanensis J5-3 T Devosiaceae Devosiaceae
Devosia riboflavina IFO 13584 T Devosiaceae Devosiaceae
Cohaesibacter gelatinilyticus DSM 18289 T Cohaesibacteraceae Cohaesibacteraceae
Consotaella salsifontis USBA 369 T Aurantimonadaceae Aurantimonadaceae
Mangrovicella endophytica 5T4P-12-1 T Aurantimonadaceae Aurantimonadaceae
Aureimonas altamirensis DSM 21988 T Aurantimonadaceae Aurantimonadaceae
Fulvimarina pelagi HTCC2506 T Aurantimonadaceae Aurantimonadaceae
Aurantimonas coralicida DSM 14790 T Aurantimonadaceae Aurantimonadaceae
Salaquimonas pukyongi RR3-28 T Phyllobacteriaceae Salaquimonadaceae
Oricola cellulosilytica JCM 19534 T Phyllobacteriaceae Ahrensiaceae
Roseitalea porphyridii MA7-20 T Phyllobacteriaceae Ahrensiaceae
Oceaniradius stylonematis StC1 T Phyllobacteriaceae Ahrensiaceae
Ahrensia kielensis DSM 5890 T Ahrensiaceae Ahrensiaceae
Zhengella mangrovi X9-2-2 T Notoacmeibacteraceae Bartonellaceae
Notoacmeibacter marinus XMTR2A4 T Notoacmeibacteraceae Bartonellaceae
Bartonella bacilliformis KC583 T Bartonellaceae Bartonellaceae
Phyllobacterium myrsinacearum DSM 5892 T Phyllobacteriaceae Bartonellaceae
"Paramesorhizobium deserti" A-3-E T Phyllobacteriaceae Bartonellaceae
Pseudochrobactrum asaccharolyticum DSM 25619 T Brucellaceae Bartonellaceae
Paenochrobactrum gallinarii DSM 22336 T Brucellaceae Bartonellaceae
Falsochrobactrum ovis DSM 26720 T Brucellaceae Bartonellaceae
Ochrobactrum anthropi ATCC 49188 T Brucellaceae Bartonellaceae
Brucella melitensis 16M T Brucellaceae Bartonellaceae
Limoniibacter endophyticus KCTC 42097 T Hyphomicrobiaceae Bartonellaceae
Nitratireductor aquibiodomus JCM 21793 T Phyllobacteriaceae Bartonellaceae
Chelativorans multitrophicus DSM 9103 T Phyllobacteriaceae Bartonellaceae
Aquibium oceanicum MCCC 1K02305 T Phyllobacteriaceae Bartonellaceae
Tianweitania sediminis Z8 T Phyllobacteriaceae Bartonellaceae
Corticibacterium populi KCTC 42249 T Phyllobacteriaceae Bartonellaceae
Pseudaminobacter salicylatoxidans DSM 6986 T Phyllobacteriaceae Bartonellaceae
Aquamicrobium defluvii DSM 11603 T Phyllobacteriaceae Bartonellaceae
Mesorhizobium loti DSM 2626 T Phyllobacteriaceae Bartonellaceae
"Oryzicola mucosus" ROOL2 T Phyllobacteriaceae Bartonellaceae
Aminobacter aminovorans DSM 7048 T Phyllobacteriaceae Bartonellaceae
"Neopararhizobium haloflavum" XC0140 T Rhizobiaceae Rhizobiaceae
Georhizobium profundi WS11 T Rhizobiaceae Rhizobiaceae
"Pararhizobium mangrovi" BGMRC 6574 T Rhizobiaceae Rhizobiaceae
Pseudohoeflea suaedae YC6898 T Phyllobacteriaceae Rhizobiaceae
Hoeflea marina DSM 16791 T Rhizobiaceae Rhizobiaceae
Onobrychidicola muellerharveyae TH2 T Rhizobiaceae Rhizobiaceae
Shinella granuli DSM 18401 T Rhizobiaceae Rhizobiaceae
Mycoplana dimorpha DSM 7138 T Rhizobiaceae Rhizobiaceae
Xaviernesmea oryzae CGMCC 1.7048 T Rhizobiaceae Rhizobiaceae
Pararhizobium giardinii H152 T Rhizobiaceae Rhizobiaceae
Sinorhizobium fredii USDA 205 T Rhizobiaceae Rhizobiaceae
Ensifer adhaerens ATCC 33212 T Rhizobiaceae Rhizobiaceae
Rhizobium leguminosarum USDA 2370 T Rhizobiaceae Rhizobiaceae
Gellertiella hungarica DSM 29853 T Rhizobiaceae Rhizobiaceae
Martelella mediterranea DSM 17316 T Rhizobiaceae Rhizobiaceae
Liberibacter crescens BT-1 T Rhizobiaceae Rhizobiaceae
Pseudorhizobium marinum MGL06 T Rhizobiaceae Rhizobiaceae
Neorhizobium galegae HAMBI 540 T Rhizobiaceae Rhizobiaceae
Endobacterium cereale RZME27 T Rhizobiaceae Rhizobiaceae
Agrobacterium tumefaciens B6 T Rhizobiaceae Rhizobiaceae
Peteryoungia ipomoeae shin9-1 T Rhizobiaceae Rhizobiaceae
Ciceribacter lividus DSM 25528 T Rhizobiaceae Rhizobiaceae
Allorhizobium undicola ATCC 700741 T Rhizobiaceae Rhizobiaceae
Faunimonas pinastri A52C2 T Pleomorphomonadaceae Afifellaceae
Afifella marina DSM 2698 T Afifellaceae Afifellaceae
Prosthecomicrobium pneumaticum DSM 16268 T Kaistiaceae Kaistiaceae
Kaistia adipata DSM 17808 T Kaistiaceae Kaistiaceae
Bauldia litoralis ATCC 35022 T Kaistiaceae Kaistiaceae
Methylobrevis pamukkalensis VKM B-2849 T Pleomorphomonadaceae Pleomorphomonadaceae
Hartmannibacter diazotrophicus E19 T Pleomorphomonadaceae Pleomorphomonadaceae
Pleomorphomonas oryzae DSM 16300 T Pleomorphomonadaceae Pleomorphomonadaceae
Oharaeibacter diazotrophicus DSM 102969 T Pleomorphomonadaceae Pleomorphomonadaceae
Mongoliimonas terrestris MIMtkB18 T Pleomorphomonadaceae Pleomorphomonadaceae
Chthonobacter albigriseus KCTC 42450 T Pleomorphomonadaceae Pleomorphomonadaceae
Siculibacillus lacustris SA-279 T Ancalomicrobiaceae Ancalomicrobiaceae
Rhodobium orientis DSM 11290 T Rhodobiaceae Rhodobiaceae
Pseudovibrio denitrificans DSM 17465 T Stappiaceae Stappiaceae
Roseibium denhamense JCM 10543 T Stappiaceae Stappiaceae
Pannonibacter phragmitetus DSM 14782 T Stappiaceae Stappiaceae
Stappia stellulata DSM 5886 T Stappiaceae Stappiaceae
Hongsoonwoonella zoysiae SY4-7 T Stappiaceae Stappiaceae
Breoghania corrubedonensis DSM 23382 T Breoghaniaceae Stappiaceae
Segnochrobactrum spirostomi Sp-1 T Segnochrobactraceae Segnochrobactraceae
Pseudoxanthobacter soli DSM 19599 T Pseudoxanthobacteraceae Segnochrobactraceae
Amorphus coralli DSM 19760 T Amorphaceae Amorphaceae
Acuticoccus yangtzensis JL1095 T Amorphaceae Amorphaceae
Tree scale: 0.1

**Figure S2. Phylogenetic analysis of the order *Hyphomicrobiales*.** On the left, a maximum likelihood phylogeny of 138 *Hyphomicrobiales* type strains is shown, built using the concatenated protein alignments encoded by the core_143 gene set (19 genes present in 100% of the strains). The phylogeny was rooted using five *Caulobacterales* type strains as the outgroup. The numbers on the nodes indicate the ultra-fast jackknife values using a 40% resampling rate (top numbers) and the SH-aLRT support values (bottom numbers), both calculated from 1000 replicates. Only values below 100 are shown. The scale bar represents the average number of amino acid substitutions per site. To the right of the phylogeny is the current family assignments of each of the 138 *Hyphomicrobiales* type strains, followed to the right by the proposed family assignments of each strain.

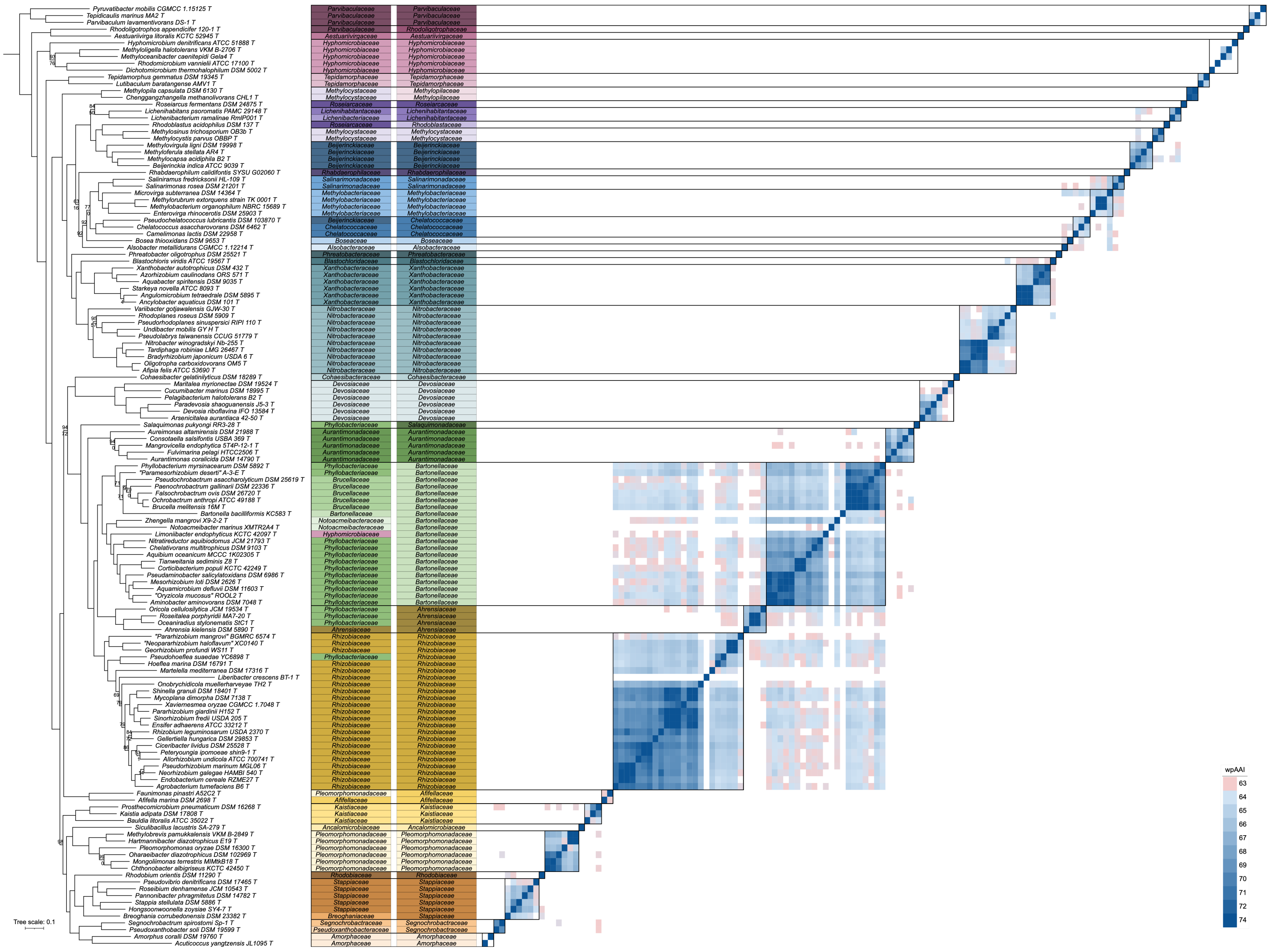
Pyruvatibacter mobilis CGMCC 1.15125 T
Tepidicaulis marinus MA2 T
Parvibaculum lavamentivorans DS-1 T
Rhodoligotrophos appendicifer 120-1 T
Aestuariivirga litoralis KCTC 52945 T
Hyphomicrobium denitrificans ATCC 51888 T
Methyloligella halotolerans VKM B-2706 T
Methyloceanibacter caenitepidi Gela4 T
Rhodomicrobium vannielii ATCC 17100 T
Dichotomicrobium thermohalophilum DSM 5002 T
Tepidamorphus gemmatus DSM 19345 T
Lutibaculum baratangense AMV1 T
Methylopila capsulata DSM 6130 T
Chenggangzhangella methanolivorans CHL1 T
Roseiarcus fermentans DSM 24875 T
Lichenihabitans psoromatis PAMC 29148 T
Lichenibacterium ramalinae RmlP001 T
Rhodoblastus acidophilus DSM 137 T
Methylosinus trichosporium OB3b T
Methylocystis parvus OBBP T
Methylovirgula ligni DSM 19998 T
Methyloferula stellata AR4 T
Methylocapsa acidiphila B2 T
Beijerinckia indica ATCC 9039 T
Rhabdaerophilum calidifontis SYSU G02060 T
Saliniramus fredricksonii HL-109 T
Salinarimonas rosea DSM 21201 T
Microvirga subterranea DSM 14364 T
Methylorubrum extorquens strain TK 0001 T
Methylobacterium organophilum NBRC 15689 T
Enterovirga rhinocerotis DSM 25903 T
Pseudochelatococcus lubricantis DSM 103870 T
Chelatococcus asaccharovorans DSM 6462 T
Camelimonas lactis DSM 22958 T
Bosea thioxidans DSM 9653 T
Alsobacter metallidurans CGMCC 1.12214 T
Phreatobacter oligotrophus DSM 25521 T
Blastochloris viridis ATCC 19567 T
Xanthobacter autotrophicus DSM 432 T
Azorhizobium caulinodans ORS 571 T
Aquabacter spiritensis DSM 9035 T
Starkeya novella ATCC 8093 T
Angulomicrobium tetraedrale DSM 5895 T
Ancylobacter aquaticus DSM 101 T
Variibacter gotjawalensis GJW-30 T
Rhodoplanes roseus DSM 5909 T
Pseudorhodoplanes sinuspersici RIPI 110 T
Undibacter mobilis GY H T
Pseudolabrys taiwanensis CCUG 51779 T
Nitrobacter winogradskyi Nb-255 T
Tardiphaga robiniae LMG 26467 T
Bradyrhizobium japonicum USDA 6 T
Oligotropha carboxidovorans OM5 T
Afipia felis ATCC 53690 T
Cohaesibacter gelatinilyticus DSM 18289 T
Maritalea myrionectae DSM 19524 T
Cucumibacter marinus DSM 18995 T
Pelagibacterium halotolerans B2 T
Paradevosia shaoguanensis J5-3 T
Devosia riboflavina IFO 13584 T
Arsenicitalea aurantiaca 42-50 T
Salaquimonas pukyongi RR3-28 T
Aureimonas altamirensis DSM 21988 T
Consotaella salsifontis USBA 369 T
Mangrovicella endophytica ST4P-12-1 T
Fulvimarina pelagi HTCC2506 T
Aurantimonas coralicida DSM 14790 T
Phyllobacterium myrsinacearum DSM 5892 T
"Paramesorhizobium deserti" A-3-E T
Pseudochrobactrum asaccharolyticum DSM 25619 T
Paenochrobactrum gallinarii DSM 22336 T
Falsochrobactrum ovis DSM 26720 T
Ochrobactrum anthropi ATCC 49188 T
Brucella melitensis 16M T
Bartonella bacilliformis KC583 T
Zhengella mangrovi X9-2-2 T
Notoacmeibacter marinus XMTR2A4 T
Limoniibacter endophyticus KCTC 42097 T
Nitratireductor aquibiodomus JCM 21793 T
Chelativorans multitrophicus DSM 9103 T
Aquibium oceanicum MCCC 1K02305 T
Tianweitania sediminis Z8 T
Corticibacterium populi KCTC 42249 T
Pseudaminobacter salicylatoxidans DSM 6986 T
Mesorhizobium loti DSM 2626 T
Aquamicrobium defluvii DSM 11603 T
"Oryzicola mucosus" ROOL2 T
Aminobacter aminovorans DSM 7048 T
Oricola cellulosilytica JCM 19534 T
Roseitalea porphyridii MA7-20 T
Oceaniradius stylonematis StC1 T
Ahrensia kielensis DSM 5890 T
"Pararhizobium mangrovi" BGMRC 6574 T
"Neopararhizobium haloflavum" XC0140 T
Georhizobium profundi WS11 T
Pseudohoeflea suaedae YC6898 T
Hoeflea marina DSM 16791 T
Martelella mediterranea DSM 17316 T
Liberibacter crescens BT-1 T
Onobrychidicola muellerharveyae TH2 T
Shinella granuli DSM 18401 T
Mycoplana dimorpha DSM 7138 T
Xaviernesmea oryzae CGMCC 1.7048 T
Pararhizobium giardinii H152 T
Sinorhizobium fredii USDA 205 T
Ensifer adhaerens ATCC 33212 T
Rhizobium leguminosarum USDA 2370 T
Gellertiella hungarica DSM 29853 T
Ciceribacter lividus DSM 25528 T
Peteryoungia ipomoeae shin9-1 T
Allorhizobium undicola ATCC 700741 T
Pseudorhizobium marinum MGL06 T
Neorhizobium galegae HAMBI 540 T
Endobacterium cereale RZME27 T
Agrobacterium tumefaciens B6 T
Faunimonas pinastri A52C2 T
Afifella marina DSM 2698 T
Prosthecomicrobium pneumaticum DSM 16268 T
Kaistia adipata DSM 17808 T
Bauldia litoralis ATCC 35022 T
Siculibacillus lacustris SA-279 T
Methylobrevis pamukkalensis VKM B-2849 T
Hartmannibacter diazotrophicus E19 T
Pleomorphomonas oryzae DSM 16300 T
Oharaeibacter diazotrophicus DSM 102969 T
Mongoliimonas terrestris MIMtkB18 T
Chthonobacter albigriseus KCTC 42450 T
Rhodobium orientis DSM 11290 T
Pseudovibrio denitrificans DSM 17465 T
Roseibium denhamense JCM 10543 T
Pannonibacter phragmitetus DSM 14782 T
Stappia stellulata DSM 5886 T
Hongsoonwoonella zoysiae SY4-7 T
Breoghania corrubedonensis DSM 23382 T
Segnochrobactrum spirostomi Sp-1 T
Pseudoxanthobacter soli DSM 19599 T
Amorphus coralli DSM 19760 T
Acuticoccus yangtzensis JL1095 T
Tree scale: 0.1
wpAAI
74

**Figure S3. Phylogenetic and whole-proteome AAI (wpAA) analyses of the order *Hyphomicrobiales*.** On the left, a maximum likelihood phylogeny of 138 *Hyphomicrobiales* type strains is shown, built using the concatenated protein alignments encoded by the perc95_143 gene set (256 genes present in at least 95% of the strains). The phylogeny was rooted using five *Caulobacterales* type strains as the outgroup. The numbers on the nodes indicate the ultra-fast jackknife values using a 40% resampling rate (top numbers) and the SH-aLRT support values (bottom numbers), both calculated from 1000 replicates. Only values below 100 are shown. The scale bar represents the average number of amino acid substitutions per site. To the right of the phylogeny is the current family assignments of each of the 138 *Hyphomicrobiales* type strains, followed to the right by the proposed family assignments of each strain. On the righthand side, a matrix is provided showing the wpAAI values between each pair of strains calculated using EzAAI. Values less than 63% are in white while all values greater than 73% are the same shade of blue. Black boxes indicate the proposed families.

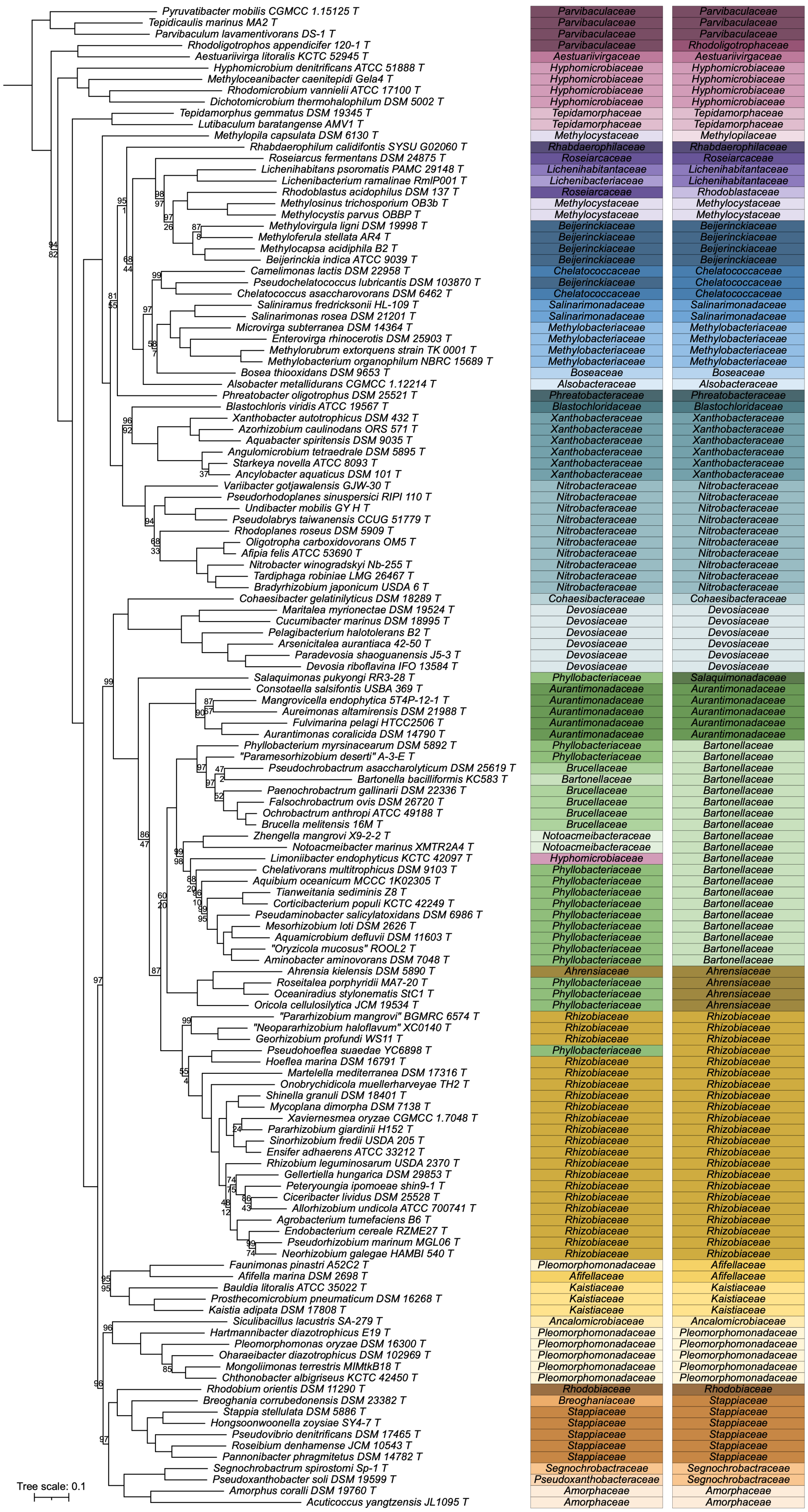
Pyruvatibacter mobilis CGMCC 1.15125 T Parvibaculaceae Parvibaculaceae
Tepidicaulis marinus MA2 T Parvibaculaceae Parvibaculaceae
Parvibaculum lavamentivorans DS-1 T Parvibaculaceae Parvibaculaceae
Rhodoligotrophos appendicifer 120-1 T Parvibaculaceae Rhodoligotrophaceae
Aestuariivirga litoralis KCTC 52945 T Aestuariivirgaceae Aestuariivirgaceae
Hyphomicrobium denitrificans ATCC 51888 T Hyphomicrobiaceae Hyphomicrobiaceae
Methyloceanibacter caenitepidi Gela4 T Hyphomicrobiaceae Hyphomicrobiaceae
Rhodomicrobium vannielii ATCC 17100 T Hyphomicrobiaceae Hyphomicrobiaceae
Dichotomicrobium thermohalophilum DSM 5002 T Hyphomicrobiaceae Hyphomicrobiaceae
Tepidamorphus gemmatus DSM 19345 T Tepidamorphaceae Tepidamorphaceae
Lutibaculum baratangense AMV1 T Tepidamorphaceae Tepidamorphaceae
Methylopila capsulata DSM 6130 T Methylocystaceae Methylopilaceae
Rhabdaerophilum calidifontis SYSU G02060 T Rhabdaerophilaceae Rhabdaerophilaceae
Roseiarcus fermentans DSM 24875 T Roseiarcaceae Roseiarcaceae
Lichenihabitans psoromatis PAMC 29148 T Lichenihabitantaceae Lichenihabitantaceae
Lichenibacterium ramalinae RmlP001 T Lichenibacteriaceae Lichenihabitantaceae
Rhodoblastus acidophilus DSM 137 T Roseiarcaceae Rhodoblastaceae
Methylosinus trichosporium OB3b T Methylocystaceae Methylocystaceae
Methylocystis parvus OBBP T Methylocystaceae Methylocystaceae
Methylovirgula ligni DSM 19998 T Beijerinckiaceae Beijerinckiaceae
Methyloferula stellata AR4 T Beijerinckiaceae Beijerinckiaceae
Methylocapsa acidiphila B2 T Beijerinckiaceae Beijerinckiaceae
Beijerinckia indica ATCC 9039 T Beijerinckiaceae Beijerinckiaceae
Camelimonas lactis DSM 22958 T Chelatococcaceae Chelatococcaceae
Pseudochelatococcus lubricantis DSM 103870 T Beijerinckiaceae Chelatococcaceae
Chelatococcus asaccharovorans DSM 6462 T Chelatococcaceae Chelatococcaceae
Saliniramus fredricksonii HL-109 T Salinarimonadaceae Salinarimonadaceae
Salinarimonas rosea DSM 21201 T Salinarimonadaceae Salinarimonadaceae
Microvirga subterranea DSM 14364 T Methylobacteriaceae Methylobacteriaceae
Enterovirga rhinocerotis DSM 25903 T Methylobacteriaceae Methylobacteriaceae
Methylorubrum extorquens strain TK 0001 T Methylobacteriaceae Methylobacteriaceae
Methylobacterium organophilum NBRC 15689 T Methylobacteriaceae Methylobacteriaceae
Bosea thiooxidans DSM 9653 T Boseaceae Boseaceae
Alsobacter metallidurans CGMCC 1.12214 T Alsobacteraceae Alsobacteraceae
Phreatobacter oligotrophus DSM 25521 T Phreatobacteraceae Phreatobacteraceae
Blastochloris viridis ATCC 19567 T Blastochloridaceae Blastochloridaceae
Xanthobacter autotrophicus DSM 432 T Xanthobacteraceae Xanthobacteraceae
Azorhizobium caulinodans ORS 571 T Xanthobacteraceae Xanthobacteraceae
Aquabacter spiritensis DSM 9035 T Xanthobacteraceae Xanthobacteraceae
Angulomicrobium tetraedrale DSM 5895 T Xanthobacteraceae Xanthobacteraceae
Starkeya novella ATCC 8093 T Xanthobacteraceae Xanthobacteraceae
Ancylobacter aquaticus DSM 101 T Xanthobacteraceae Xanthobacteraceae
Variibacter gotjawalensis GJW-30 T Nitrobacteraceae Nitrobacteraceae
Pseudorhodoplanes sinuspersici RIPI 110 T Nitrobacteraceae Nitrobacteraceae
Undibacter mobilis GY H T Nitrobacteraceae Nitrobacteraceae
Pseudolabrys taiwanensis CCUG 51779 T Nitrobacteraceae Nitrobacteraceae
Rhodoplanes roseus DSM 5909 T Nitrobacteraceae Nitrobacteraceae
Oligotropha carboxidovorans OM5 T Nitrobacteraceae Nitrobacteraceae
Afipia felis ATCC 53690 T Nitrobacteraceae Nitrobacteraceae
Nitrobacter winogradskyi Nb-255 T Nitrobacteraceae Nitrobacteraceae
Tardiphaga robiniae LMG 26467 T Nitrobacteraceae Nitrobacteraceae
Bradyrhizobium japonicum USDA 6 T Nitrobacteraceae Nitrobacteraceae
Cohaesibacter gelatinilyticus DSM 18289 T Cohaesibacteraceae Cohaesibacteraceae
Maritalea myrionectae DSM 19524 T Devosiaceae Devosiaceae
Cucumibacter marinus DSM 18995 T Devosiaceae Devosiaceae
Pelagibacterium halotolerans B2 T Devosiaceae Devosiaceae
Arsenicitalea aurantiaca 42-50 T Devosiaceae Devosiaceae
Paradevosia shaoguanensis J5-3 T Devosiaceae Devosiaceae
Devosia riboflavina IFO 13584 T Devosiaceae Devosiaceae
Salaquimonas pukyongi RR3-28 T Phyllobacteriaceae Salaquimonadaceae
Consotaella salsifontis USBA 369 T Aurantimonadaceae Aurantimonadaceae
Mangrovicella endophytica 5T4P-12-1 T Aurantimonadaceae Aurantimonadaceae
Aureimonas altamirensis DSM 21988 T Aurantimonadaceae Aurantimonadaceae
Fulvimarina pelagi HTCC2506 T Aurantimonadaceae Aurantimonadaceae
Aurantimonas coralicida DSM 14790 T Aurantimonadaceae Aurantimonadaceae
Phyllobacterium myrsinacearum DSM 5892 T Phyllobacteriaceae Bartonellaceae
"Paramesorhizobium deserti" A-3-E T Phyllobacteriaceae Bartonellaceae
Pseudochrobactrum asaccharolyticum DSM 25619 T Brucellaceae Bartonellaceae
Bartonella bacilliformis KC583 T Bartonellaceae Bartonellaceae
Paenochrobactrum gallinarii DSM 22336 T Brucellaceae Bartonellaceae
Falsochrobactrum ovis DSM 26720 T Brucellaceae Bartonellaceae
Ochrobactrum anthropi ATCC 49188 T Brucellaceae Bartonellaceae
Brucella melitensis 16M T Brucellaceae Bartonellaceae
Zhengella mangrovi X9-2-2 T Notoacmeibacteraceae Bartonellaceae
Notoacmeibacter marinus XMTR2A4 T Notoacmeibacteraceae Bartonellaceae
Limoniibacter endophyticus KCTC 42097 T Hyphomicrobiaceae Bartonellaceae
Chelativorans multitrophicus DSM 9103 T Phyllobacteriaceae Bartonellaceae
Aquibium oceanicum MCCC 1K02305 T Phyllobacteriaceae Bartonellaceae
Tianweitania sediminis Z8 T Phyllobacteriaceae Bartonellaceae
Corticibacterium populi KCTC 42249 T Phyllobacteriaceae Bartonellaceae
Pseudaminobacter salicylatoxidans DSM 6986 T Phyllobacteriaceae Bartonellaceae
Mesorhizobium loti DSM 2626 T Phyllobacteriaceae Bartonellaceae
Aquamicrobium defluvii DSM 11603 T Phyllobacteriaceae Bartonellaceae
"Oryzicola mucosus" ROOL2 T Phyllobacteriaceae Bartonellaceae
Aminobacter aminovorans DSM 7048 T Phyllobacteriaceae Bartonellaceae
Ahrensia kielensis DSM 5890 T Ahrensiaceae Ahrensiaceae
Roseitalea porphyridii MA7-20 T Phyllobacteriaceae Ahrensiaceae
Oceaniradius stylonematis StC1 T Phyllobacteriaceae Ahrensiaceae
Oricola cellulosilytica JCM 19534 T Phyllobacteriaceae Ahrensiaceae
"Pararhizobium mangrovi" BGMRC 6574 T Rhizobiaceae Rhizobiaceae
"Neopararhizobium haloflavum" XC0140 T Rhizobiaceae Rhizobiaceae
Georhizobium profundi WS11 T Rhizobiaceae Rhizobiaceae
Pseudohoeflea suaedae YC6898 T Phyllobacteriaceae Rhizobiaceae
Hoeflea marina DSM 16791 T Rhizobiaceae Rhizobiaceae
Martelella mediterranea DSM 17316 T Rhizobiaceae Rhizobiaceae
Onobrychidicola muellerharveyae TH2 T Rhizobiaceae Rhizobiaceae
Shinella granuli DSM 18401 T Rhizobiaceae Rhizobiaceae
Mycoplana dimorpha DSM 7138 T Rhizobiaceae Rhizobiaceae
Xaviernesmea oryzae CGMCC 1.7048 T Rhizobiaceae Rhizobiaceae
Pararhizobium giardinii H152 T Rhizobiaceae Rhizobiaceae
Sinorhizobium fredii USDA 205 T Rhizobiaceae Rhizobiaceae
Ensifer adhaerens ATCC 33212 T Rhizobiaceae Rhizobiaceae
Rhizobium leguminosarum USDA 2370 T Rhizobiaceae Rhizobiaceae
Gellertiella hungarica DSM 29853 T Rhizobiaceae Rhizobiaceae
Peteryoungia ipomoeae shin9-1 T Rhizobiaceae Rhizobiaceae
Ciceribacter lividus DSM 25528 T Rhizobiaceae Rhizobiaceae
Allorhizobium undicola ATCC 700741 T Rhizobiaceae Rhizobiaceae
Agrobacterium tumefaciens B6 T Rhizobiaceae Rhizobiaceae
Endobacterium cereale RZME27 T Rhizobiaceae Rhizobiaceae
Pseudorhizobium marinum MGL06 T Rhizobiaceae Rhizobiaceae
Neorhizobium galegae HAMBI 540 T Rhizobiaceae Rhizobiaceae
Faunimonas pinastri A52C2 T Pleomorphomonadaceae Afifellaceae
Afifella marina DSM 2698 T Afifellaceae Afifellaceae
Bauldia litoralis ATCC 35022 T Kaistiaceae Kaistiaceae
Prosthecomicrobium pneumaticum DSM 16268 T Kaistiaceae Kaistiaceae
Kaistia adipata DSM 17808 T Kaistiaceae Kaistiaceae
Siculibacillus lacustris SA-279 T Ancalomicrobiaceae Ancalomicrobiaceae
Hartmannibacter diazotrophicus E19 T Pleomorphomonadaceae Pleomorphomonadaceae
Pleomorphomonas oryzae DSM 16300 T Pleomorphomonadaceae Pleomorphomonadaceae
Oharaeibacter diazotrophicus DSM 102969 T Pleomorphomonadaceae Pleomorphomonadaceae
Mongoliimonas terrestris MIMtkB18 T Pleomorphomonadaceae Pleomorphomonadaceae
Chthonobacter albigriseus KCTC 42450 T Pleomorphomonadaceae Pleomorphomonadaceae
Rhodobium orientis DSM 11290 T Rhodobiaceae Rhodobiaceae
Breoghania corrubedonensis DSM 23382 T Breoghaniaceae Stappiaceae
Stappia stellulata DSM 5886 T Stappiaceae Stappiaceae
Hongsoonwoonella zoysiae SY4-7 T Stappiaceae Stappiaceae
Pseudovibrio denitrificans DSM 17465 T Stappiaceae Stappiaceae
Roseibium denhamense JCM 10543 T Stappiaceae Stappiaceae
Pannonibacter phragmitetus DSM 14782 T Stappiaceae Stappiaceae
Segnochrobactrum spirostomi Sp-1 T Segnochrobactraceae Segnochrobactraceae
Pseudoxanthobacter soli DSM 19599 T Pseudoxanthobacteraceae Segnochrobactraceae
Amorphus coralli DSM 19760 T Amorphaceae Amorphaceae
Acuticoccus yangtzensis JL1095 T Amorphaceae Amorphaceae
Tree scale: 0.1

**Figure S4. Phylogenetic analysis of the order *Hyphomicrobiales*.** On the left, a maximum likelihood phylogeny of 133 *Hyphomicrobiales* type strains is shown, built using the concatenated protein alignments encoded by the core_138 gene set (59 genes present in 100% of the strains). The phylogeny was rooted using five *Caulobacterales* type strains as the outgroup. The numbers on the nodes indicate the ultra-fast jackknife values using a 40% resampling rate (top numbers) and the SH-aLRT support values (bottom numbers), both calculated from 1000 replicates. Only values below 100 are shown. The scale bar represents the average number of amino acid substitutions per site. To the right of the phylogeny is the current family assignments of each of the 133 *Hyphomicrobiales* type strains, followed to the right by the proposed family assignments of each strain.

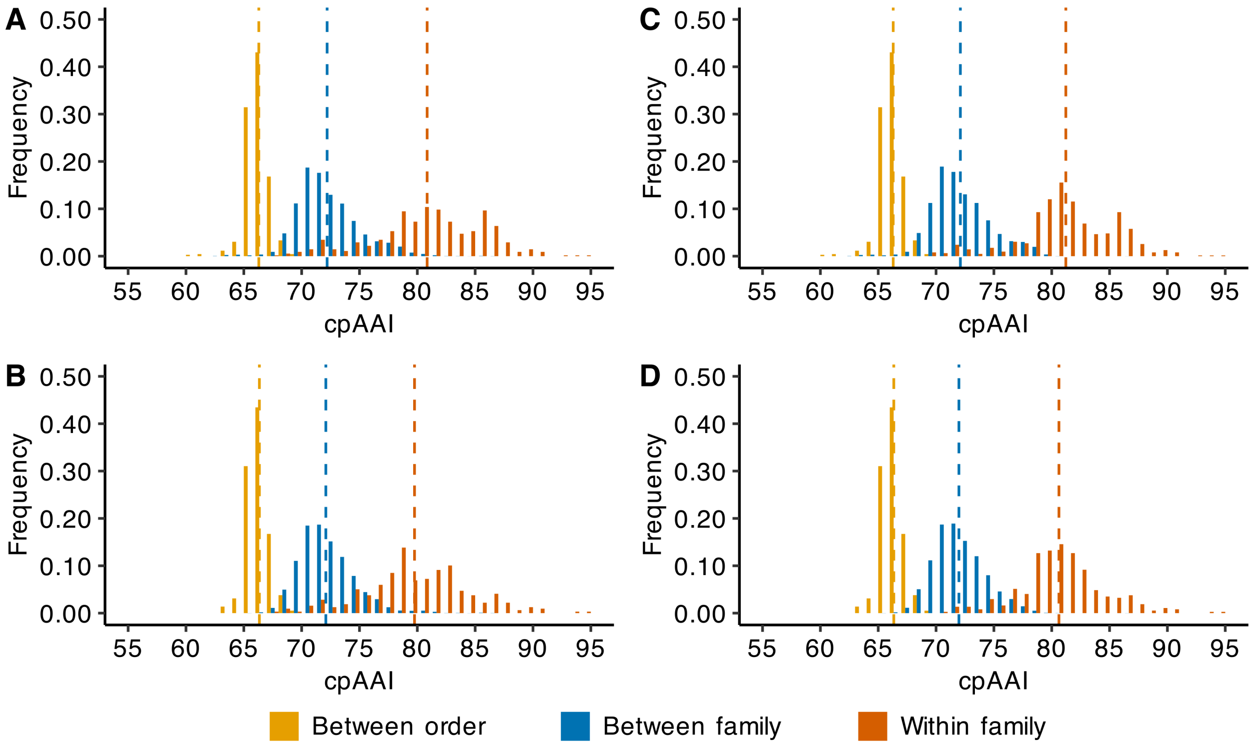

**Figure S5. Distribution of core-proteome AAI (cpAAI) comparisons of the order *Hyphomicrobiales*.** Pairwise cpAAI values were calculated based on 19 nonrecombinant loci from the core genome of 138 members of the order *Hyphomicrobiales* and five members of the order *Caulobacterales*. Results are summarized as histograms with a bin width of 1%. The cpAAI values calculated between two strains belonging to different orders (yellow), different families but same order (blue), or the same family (red) are summarized separately. Dashed vertical lines represent the mean value of each distribution. In all plots, cpAAI values where both strains belong to the order *Caulobacterales* were excluded. (**A**) The distribution of all pairwise cpAAI values with the classification (i.e., between order, between family, or within family) based on existing taxonomic assignments. (**B**) The distribution of all pairwise cpAAI values except for those including at least one strain from the family *Rhizobiaceae*, with the classification based on existing taxonomic assignments. (**C**) The distribution of all pairwise cpAAI values with the classification based on the proposed taxonomic assignments. (**D**) The distribution of all pairwise cpAAI values except for those including at least one strain from the family *Rhizobiaceae*, with the classification based on the proposed taxonomic assignments.

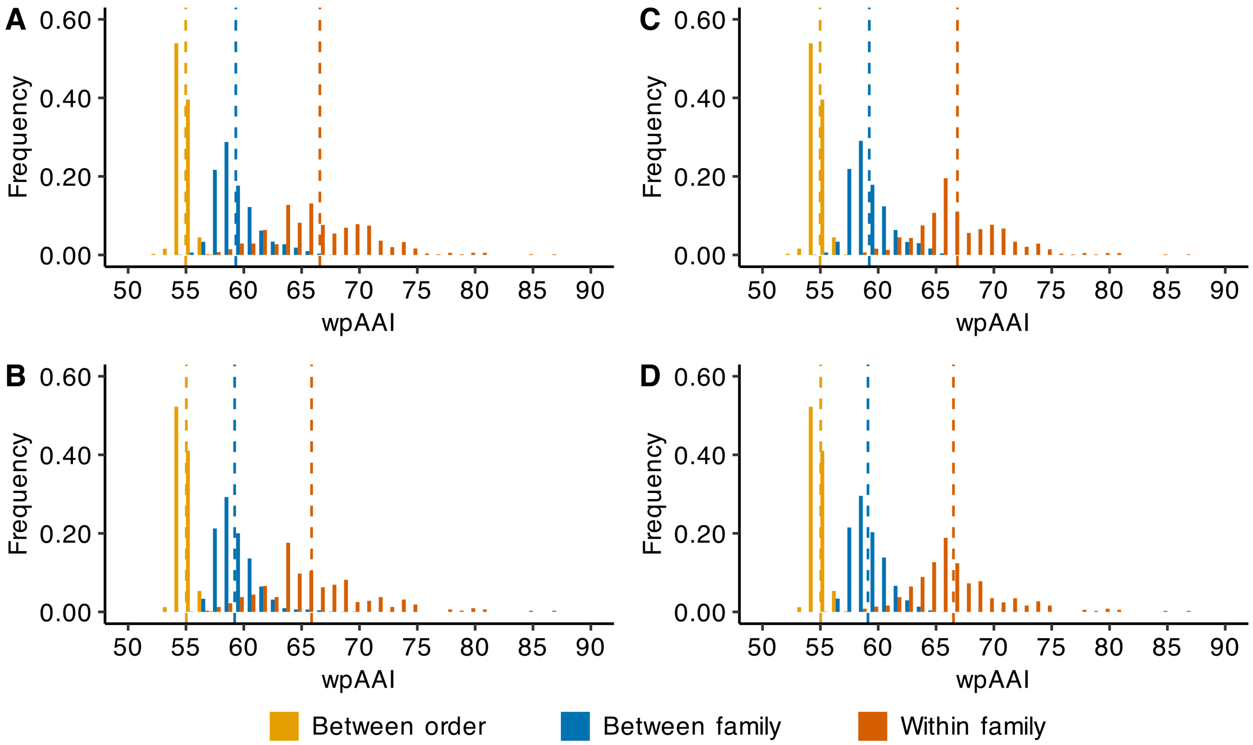

**Figure S6. Distribution of whole-proteome AAI (wpAAI) comparisons of the order *Hyphomicrobiales*.** Pairwise wpAAI values were calculated between 138 members of the order *Hyphomicrobiales* and five members of the order *Caulobacterales*. Results are summarized as histograms with a bin width of 1%. The wpAAI values calculated between two strains belonging to different orders (yellow), different families but same order (blue), or the same family (red) are summarized separately. Dashed vertical lines represent the mean value of each distribution. In all plots, wpAAI values where both strains belong to the order *Caulobacterales* were excluded. (**A**) The distribution of all pairwise wpAAI values with the classification (i.e., between order, between family, or within family) based on existing taxonomic assignments. (**B**) The distribution of all pairwise wpAAI values except for those including at least one strain from the family *Rhizobiaceae*, with the classification based on existing taxonomic assignments. (**C**) The distribution of all pairwise wpAAI values with the classification based on the proposed taxonomic assignments. (**D**) The distribution of all pairwise wpAAI values except for those including at least one strain from the family *Rhizobiaceae*, with the classification based on the proposed taxonomic assignments.

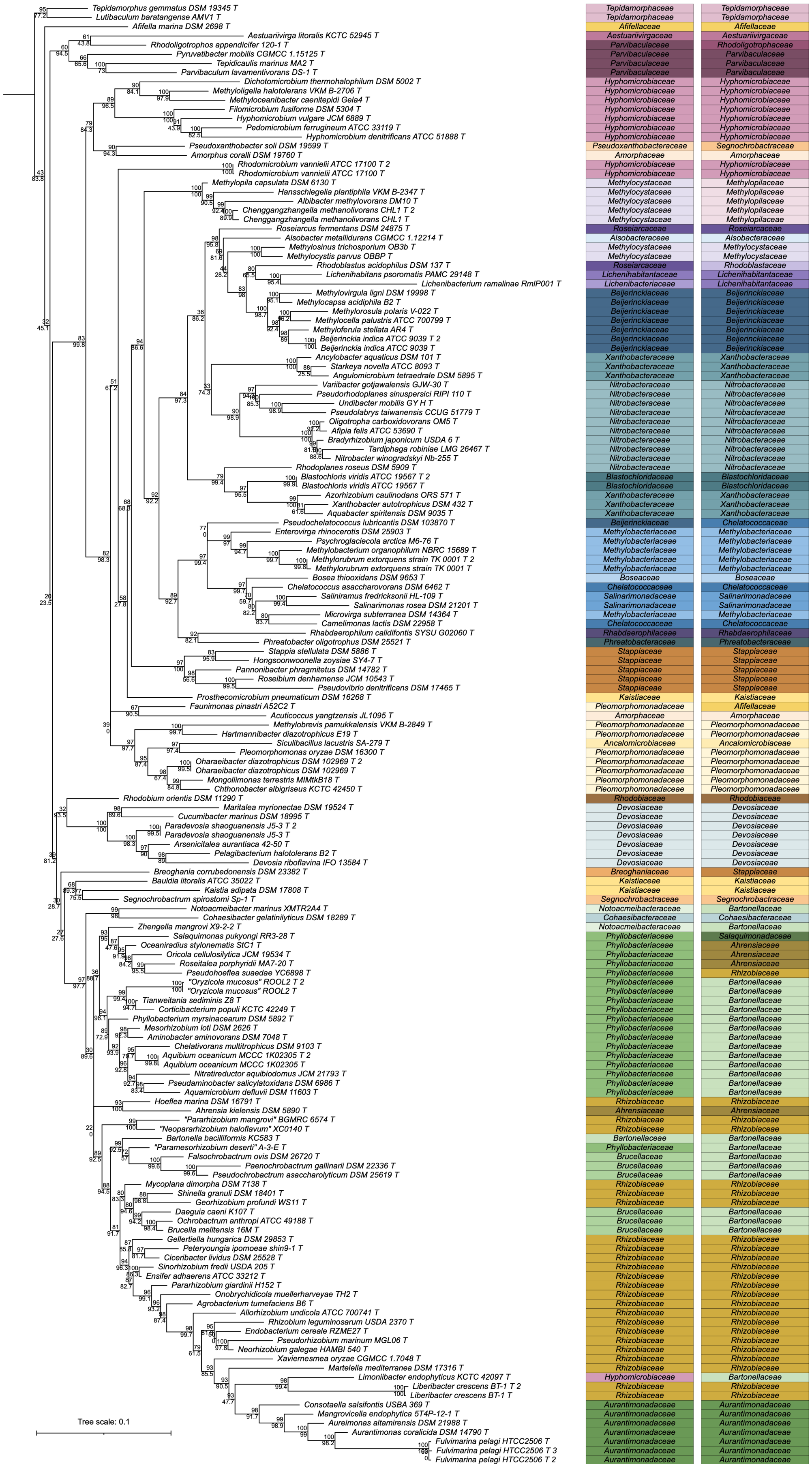
Tepidamorphus gemmatus DSM 19345 T Tepidamorphaceae Tepidamorphaceae
Lutibaculum baratangense AMV1 T Tepidamorphaceae Tepidamorphaceae
Afifella marina DSM 2698 T Afifellaceae Afifellaceae
Aestuariivirga litoralis KCTC 52945 T Aestuariivirgaceae Aestuariivirgaceae
Rhodoligotrophos appendicifer 120-1 T Parvibaculaceae Rhodoligotrophaceae
Pyruvatibacter mobilis CGMCC 1.15125 T Parvibaculaceae Parvibaculaceae
Tepidicaulis marinus MA2 T Parvibaculaceae Parvibaculaceae
Parvibaculum lavamentivorans DS-1 T Parvibaculaceae Parvibaculaceae
Dichotomicrobium thermohalophilum DSM 5002 T Hyphomicrobiaceae Hyphomicrobiaceae
Methyloligella halotolerans VKM B-2706 T Hyphomicrobiaceae Hyphomicrobiaceae
Methyloceanibacter caenitepidi Gela4 T Hyphomicrobiaceae Hyphomicrobiaceae
Filomicrobium fusiforme DSM 5304 T Hyphomicrobiaceae Hyphomicrobiaceae
Hyphomicrobium vulgare JCM 6889 T Hyphomicrobiaceae Hyphomicrobiaceae
Pedomicrobium ferrugineum ATCC 33119 T Hyphomicrobiaceae Hyphomicrobiaceae
Hyphomicrobium denitrificans ATCC 51888 T Hyphomicrobiaceae Hyphomicrobiaceae
Pseudoxanthobacter soli DSM 19599 T Pseudoxanthobacteraceae Segnochrobactraceae
Amorphus coralli DSM 19760 T Amorphaceae Amorphaceae
Rhodomicrobium vannielii ATCC 17100 T 2 Hyphomicrobiaceae Hyphomicrobiaceae
Rhodomicrobium vannielii ATCC 17100 T Hyphomicrobiaceae Hyphomicrobiaceae
Methylopila capsulata DSM 6130 T Methylocystaceae Methylopilaceae
Hansschlegelia plantiphila VKM B-2347 T Methylocystaceae Methylopilaceae
Albibacter methylovorans DM10 T Methylocystaceae Methylopilaceae
Chenggangzhangella methanolivorans CHL1 T 2 Methylocystaceae Methylopilaceae
Chenggangzhangella methanolivorans CHL1 T Methylocystaceae Methylopilaceae
Roseiarcus fermentans DSM 24875 T Roseiarcaceae Roseiarcaceae
Alsobacter metallidurans CGMCC 1.12214 T Alsobacteraceae Alsobacteraceae
Methylosinus trichosporium OB3b T Methylocystaceae Methylocystaceae
Methylocystis parvus OBBP T Methylocystaceae Methylocystaceae
Rhodoblastus acidophilus DSM 137 T Roseiarcaceae Rhodoblastaceae
Lichenihabitans psoromatis PAMC 29148 T Lichenihabitantaceae Lichenihabitantaceae
Lichenibacterium ramalinae RmlP001 T Lichenibacteriaceae Lichenihabitantaceae
Methylovirgula ligni DSM 19998 T Beijerinckiaceae Beijerinckiaceae
Methylocapsa acidiphila B2 T Beijerinckiaceae Beijerinckiaceae
Methylorosula polaris V-022 T Beijerinckiaceae Beijerinckiaceae
Methylocella palustris ATCC 700799 T Beijerinckiaceae Beijerinckiaceae
Methyloferula stellata AR4 T Beijerinckiaceae Beijerinckiaceae
Beijerinckia indica ATCC 9039 T 2 Beijerinckiaceae Beijerinckiaceae
Beijerinckia indica ATCC 9039 T Beijerinckiaceae Beijerinckiaceae
Ancylobacter aquaticus DSM 101 T Xanthobacteraceae Xanthobacteraceae
Starkeya novella ATCC 8093 T Xanthobacteraceae Xanthobacteraceae
Angulomicrobium tetraedrale DSM 5895 T Xanthobacteraceae Xanthobacteraceae
Variibacter gotjawalensis GJW-30 T Nitrobacteraceae Nitrobacteraceae
Pseudorhodoplanes sinuspersici RIPI 110 T Nitrobacteraceae Nitrobacteraceae
Undibacter mobilis GY H T Nitrobacteraceae Nitrobacteraceae
Pseudolabrys taiwanensis CCUG 51779 T Nitrobacteraceae Nitrobacteraceae
Oligotropha carboxidovorans OM5 T Nitrobacteraceae Nitrobacteraceae
Afipia felis ATCC 53690 T Nitrobacteraceae Nitrobacteraceae
Bradyrhizobium japonicum USDA 6 T Nitrobacteraceae Nitrobacteraceae
Tardiphaga robiniae LMG 26467 T Nitrobacteraceae Nitrobacteraceae
Nitrobacter winogradskyi Nb-255 T Nitrobacteraceae Nitrobacteraceae
Rhodoplanes roseus DSM 5909 T Nitrobacteraceae Nitrobacteraceae
Blastochloris viridis ATCC 19567 T 2 Blastochloridaceae Blastochloridaceae
Blastochloris viridis ATCC 19567 T Blastochloridaceae Blastochloridaceae
Azorhizobium caulinodans ORS 571 T Xanthobacteraceae Xanthobacteraceae
Xanthobacter autotrophicus DSM 432 T Xanthobacteraceae Xanthobacteraceae
Aquabacter spiritensis DSM 9035 T Xanthobacteraceae Xanthobacteraceae
Pseudochelatococcus lubricantis DSM 103870 T Beijerinckiaceae Chelatococcaceae
Enterovirga rhinocerotis DSM 25903 T Methylobacteriaceae Methylobacteriaceae
Psychroglaciecola arctica M6-76 T Methylobacteriaceae Methylobacteriaceae
Methylobacterium organophilum NBRC 15689 T Methylobacteriaceae Methylobacteriaceae
Methylorubrum extorquens strain TK 0001 T 2 Methylobacteriaceae Methylobacteriaceae
Methylorubrum extorquens strain TK 0001 T Methylobacteriaceae Methylobacteriaceae
Bosea thiooxidans DSM 9653 T Boseaceae Boseaceae
Chelatococcus asaccharovorans DSM 6462 T Chelatococcaceae Chelatococcaceae
Saliniramus fredricksonii HL-109 T Salinarimonadaceae Salinarimonadaceae
Salinarimonas rosea DSM 21201 T Salinarimonadaceae Salinarimonadaceae
Microvirga subterranea DSM 14364 T Methylobacteriaceae Methylobacteriaceae
Camelimonas lactis DSM 22958 T Chelatococcaceae Chelatococcaceae
Rhabdaerophilum calidifontis SYSU G02060 T Rhabdaerophilaceae Rhabdaerophilaceae
Phreatobacter oligotrophus DSM 25521 T Phreatobacteraceae Phreatobacteraceae
Stappia stellulata DSM 5886 T Stappiaceae Stappiaceae
Hongsoonwoonella zoysiae SY4-7 T Stappiaceae Stappiaceae
Pannonibacter phragmitetus DSM 14782 T Stappiaceae Stappiaceae
Roseibium denhamense JCM 10543 T Stappiaceae Stappiaceae
Pseudovibrio denitrificans DSM 17465 T Stappiaceae Stappiaceae
Prosthecomicrobium pneumaticum DSM 16268 T Kaistiaceae Kaistiaceae
Faunimonas pinastri A52C2 T Pleomorphomonadaceae Afifellaceae
Acuticoccus yangtzensis JL1095 T Amorphaceae Amorphaceae
Methylobrevis pamukkalensis VKM B-2849 T Pleomorphomonadaceae Pleomorphomonadaceae
Hartmannibacter diazotrophicus E19 T Pleomorphomonadaceae Pleomorphomonadaceae
Siculibacillus lacustris SA-279 T Ancalomicrobiaceae Ancalomicrobiaceae
Pleomorphomonas oryzae DSM 16300 T Pleomorphomonadaceae Pleomorphomonadaceae
Oharaeibacter diazotrophicus DSM 102969 T 2 Pleomorphomonadaceae Pleomorphomonadaceae
Oharaeibacter diazotrophicus DSM 102969 T Pleomorphomonadaceae Pleomorphomonadaceae
Mongoliimonas terrestris MIMtkB18 T Pleomorphomonadaceae Pleomorphomonadaceae
Chthonobacter albigriseus KCTC 42450 T Pleomorphomonadaceae Pleomorphomonadaceae
Rhodobium orientis DSM 11290 T Rhodobiaceae Rhodobiaceae
Maritalea myrionectae DSM 19524 T Devosiaceae Devosiaceae
Cucumibacter marinus DSM 18995 T Devosiaceae Devosiaceae
Paradevosia shaoguanensis J5-3 T 2 Devosiaceae Devosiaceae
Paradevosia shaoguanensis J5-3 T Devosiaceae Devosiaceae
Arsenicitalea aurantiaca 42-50 T Devosiaceae Devosiaceae
Pelagibacterium halotolerans B2 T Devosiaceae Devosiaceae
Devosia riboflavina IFO 13584 T Devosiaceae Devosiaceae
Breoghania corrubedonensis DSM 23382 T Breoghaniaceae Stappiaceae
Bauldia litoralis ATCC 35022 T Kaistiaceae Kaistiaceae
Kaistia adipata DSM 17808 T Kaistiaceae Kaistiaceae
Segnochrobactrum spirostomi Sp-1 T Segnochrobactraceae Segnochrobactraceae
Notoacmeibacter marinus XMTR2A4 T Notoacmeibacteraceae Bartonellaceae
Cohaesibacter gelatinilyticus DSM 18289 T Cohaesibacteraceae Cohaesibacteraceae
Zhengella mangrovi X9-2-2 T Notoacmeibacteraceae Bartonellaceae
Salaquimonas pukyongi RR3-28 T Phyllobacteriaceae Salaquimonadaceae
Oceaniradius stylonematis StC1 T Phyllobacteriaceae Ahrensiaceae
Oricola cellulosilytica JCM 19534 T Phyllobacteriaceae Ahrensiaceae
Roseitalea porphyridii MA7-20 T Phyllobacteriaceae Ahrensiaceae
Pseudohoeflea suaedae YC6898 T Phyllobacteriaceae Rhizobiaceae
"Oryzicola mucosus" ROOL2 T 2 Phyllobacteriaceae Bartonellaceae
"Oryzicola mucosus" ROOL2 T Phyllobacteriaceae Bartonellaceae
Tianweitania sediminis Z8 T Phyllobacteriaceae Bartonellaceae
Corticibacterium populi KCTC 42249 T Phyllobacteriaceae Bartonellaceae
Phyllobacterium myrsinacearum DSM 5892 T Phyllobacteriaceae Bartonellaceae
Mesorhizobium loti DSM 2626 T Phyllobacteriaceae Bartonellaceae
Aminobacter aminovorans DSM 7048 T Phyllobacteriaceae Bartonellaceae
Chelativorans multitrophicus DSM 9103 T Phyllobacteriaceae Bartonellaceae
Aquibium oceanicum MCCC 1K02305 T 2 Phyllobacteriaceae Bartonellaceae
Aquibium oceanicum MCCC 1K02305 T Phyllobacteriaceae Bartonellaceae
Nitratireductor aquibiodomus JCM 21793 T Phyllobacteriaceae Bartonellaceae
Pseudaminobacter salicylatoxidans DSM 6986 T Phyllobacteriaceae Bartonellaceae
Aquamicrobium defluvii DSM 11603 T Phyllobacteriaceae Bartonellaceae
Hoeflea marina DSM 16791 T Rhizobiaceae Rhizobiaceae
Ahrensia kielensis DSM 5890 T Ahrensiaceae Ahrensiaceae
"Pararhizobium mangrovi" BGMRC 6574 T Rhizobiaceae Rhizobiaceae
"Neoparrhizobium haloflavum" XC0140 T Rhizobiaceae Rhizobiaceae
Bartonella bacilliformis KC583 T Bartonellaceae Bartonellaceae
"Paramesorhizobium deserti" A-3-E T Phyllobacteriaceae Bartonellaceae
Falsochrobactrum ovis DSM 26720 T Brucellaceae Bartonellaceae
Paenochrobactrum gallinarii DSM 22336 T Brucellaceae Bartonellaceae
Pseudochrobactrum asaccharolyticum DSM 25619 T Brucellaceae Bartonellaceae
Mycoplana dimorpha DSM 7138 T Rhizobiaceae Rhizobiaceae
Shinella granuli DSM 18401 T Rhizobiaceae Rhizobiaceae
Georhizobium profundi WS11 T Rhizobiaceae Rhizobiaceae
Daeguia caeni K107 T Brucellaceae Bartonellaceae
Ochrobactrum anthropi ATCC 49188 T Brucellaceae Bartonellaceae
Brucella melitensis 16M T Brucellaceae Bartonellaceae
Gellertiella hungarica DSM 29853 T Rhizobiaceae Rhizobiaceae
Peteryoungia ipomoeae shin9-1 T Rhizobiaceae Rhizobiaceae
Ciceribacter lividus DSM 25528 T Rhizobiaceae Rhizobiaceae
Sinorhizobium fredii USDA 205 T Rhizobiaceae Rhizobiaceae
Ensifer adhaerens ATCC 33212 T Rhizobiaceae Rhizobiaceae
Pararhizobium giardinii H152 T Rhizobiaceae Rhizobiaceae
Onobrychidicola muellerharveyae TH2 T Rhizobiaceae Rhizobiaceae
Agrobacterium tumefaciens B6 T Rhizobiaceae Rhizobiaceae
Allorhizobium undicola ATCC 700741 T Rhizobiaceae Rhizobiaceae
Rhizobium leguminosarum USDA 2370 T Rhizobiaceae Rhizobiaceae
Endobacterium cereale RZME27 T Rhizobiaceae Rhizobiaceae
Pseudorhizobium marinum MGL06 T Rhizobiaceae Rhizobiaceae
Neorhizobium galegae HAMBI 540 T Rhizobiaceae Rhizobiaceae
Xaviernesmea oryzae CGMCC 1.7048 T Rhizobiaceae Rhizobiaceae
Martelella mediterranea DSM 17316 T Rhizobiaceae Rhizobiaceae
Limoniibacter endophyticus KCTC 42097 T Hyphomicrobiaceae Bartonellaceae
Liberibacter crescens BT-1 T 2 Rhizobiaceae Rhizobiaceae
Liberibacter crescens BT-1 T Rhizobiaceae Rhizobiaceae
Consotaella salsifontis USBA 369 T Aurantimonadaceae Aurantimonadaceae
Mangrovicella endophytica 5T4P-12-1 T Aurantimonadaceae Aurantimonadaceae
Aureimonas altamirensis DSM 21988 T Aurantimonadaceae Aurantimonadaceae
Aurantimonas coralicida DSM 14790 T Aurantimonadaceae Aurantimonadaceae
Fulvimarina pelagi HTCC2506 T Aurantimonadaceae Aurantimonadaceae
Fulvimarina pelagi HTCC2506 T 3 Aurantimonadaceae Aurantimonadaceae
Fulvimarina pelagi HTCC2506 T 2 Aurantimonadaceae Aurantimonadaceae
Tree scale: 0.1

**Figure S7. 16S rRNA gene phylogenetic analysis of the order *Hyphomicrobiales*.** On the left, a maximum likelihood phylogeny of 147 *Hyphomicrobiales* type strains is shown, built using a trimmed, Clustal Omega alignment of the 16S rRNA genes. The phylogeny was rooted using five *Caulobacterales* type strains as the outgroup. In cases where a genome sequence contained more than one 16S rRNA gene sequence, all unique 16S rRNA gene sequences were included. To the right of the phylogeny is the current family assignments of each of the 147 *Hyphomicrobiales* type strains, followed to the right by the proposed family assignments of each strain.

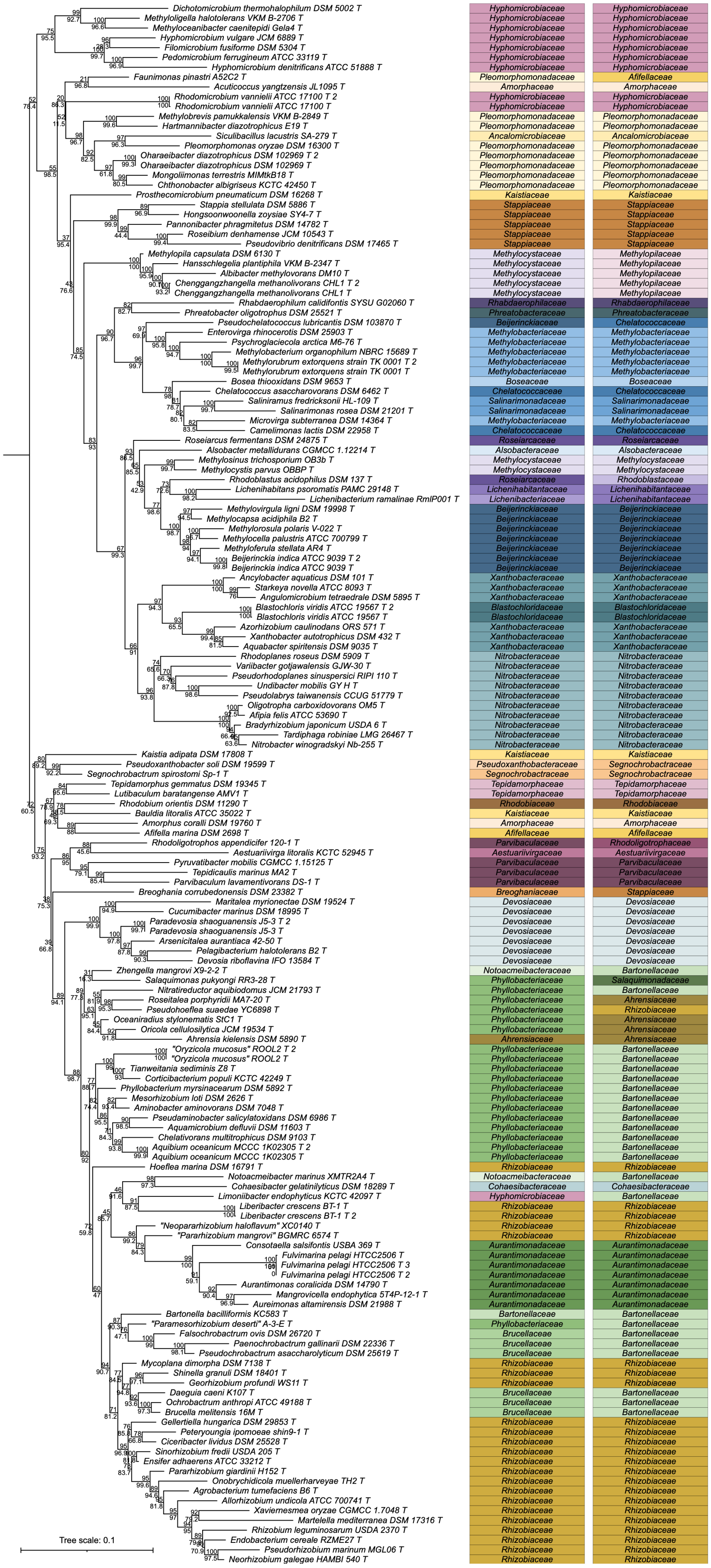

**Figure S8. 16S rRNA gene phylogenetic analysis of the order *Hyphomicrobiales*.** On the left, a maximum likelihood phylogeny of 147 *Hyphomicrobiales* type strains is shown, built using a trimmed, MAFFT alignment of the 16S rRNA genes. The phylogeny was rooted using five *Caulobacterales* type strains as the outgroup. In cases where a genome sequence contained more than one 16S rRNA gene sequence, all unique 16S rRNA gene sequences were included. To the right of the phylogeny is the current family assignments of each of the 147 *Hyphomicrobiales* type strains, followed to the right by the proposed family assignments of each strain.

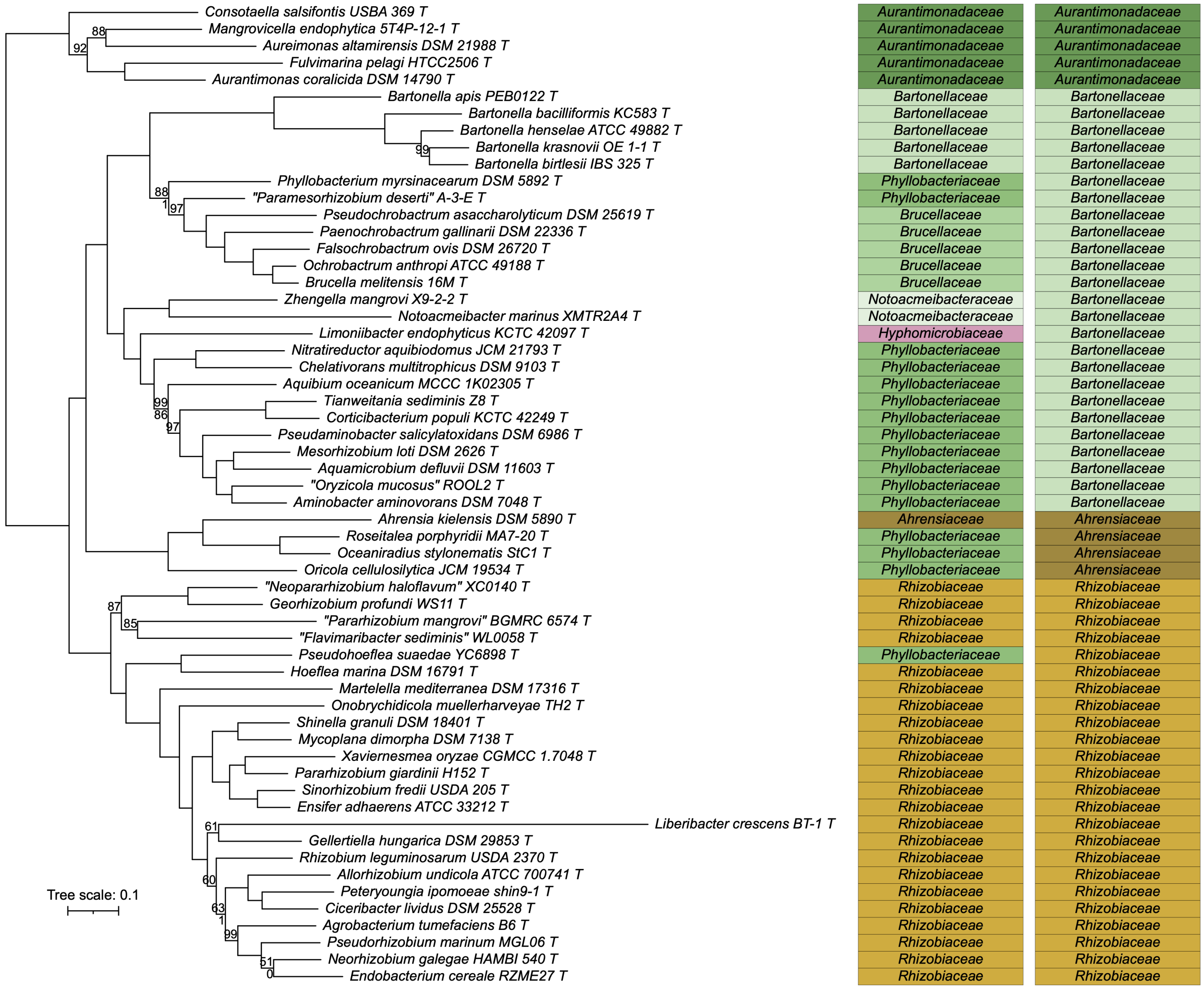

**Figure S9. Phylogenetic analysis of the family *Bartonellaceae* and related families.** On the left, an unrooted maximum likelihood phylogeny of 58 *Hyphomicrobiales* type strains is shown, built using the concatenated protein alignments encoded by the core_58 gene set (120 genes present in 100% of the included strains). The numbers on the nodes indicate the ultra-fast jackknife values using a 40% resampling rate (top numbers) and the SH-aLRT support values (bottom numbers), both calculated from 1000 replicates. Only values below 100 are shown. The scale bar represents the average number of amino acid substitutions per site. To the right of the phylogeny is the current family assignments of each of the 58 strains, followed to the right by the proposed family assignments of each strain.

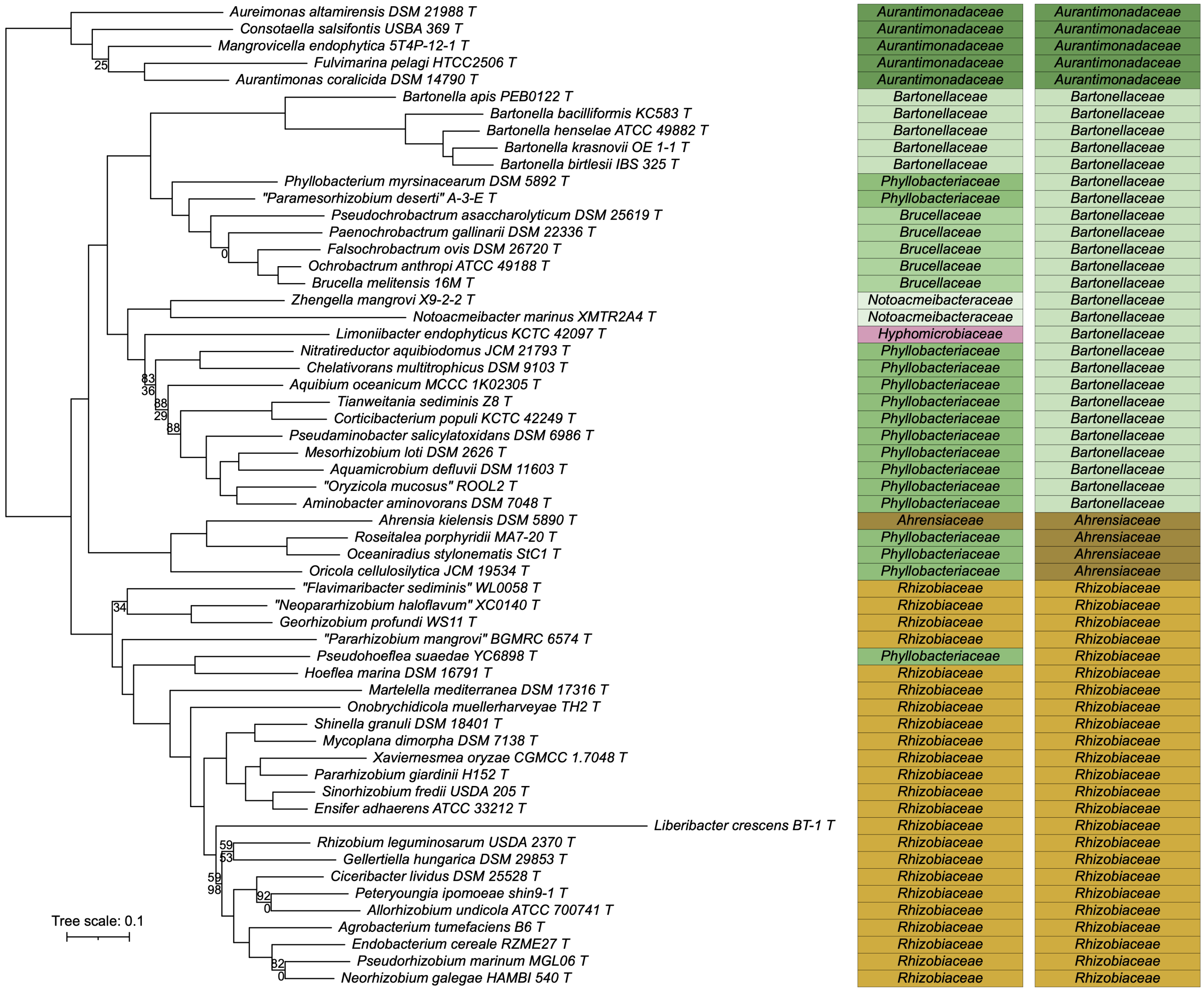

**Figure S10. Phylogenetic analysis of the family *Bartonellaceae* and related families.** On the left, an unrooted maximum likelihood phylogeny of 58 *Hyphomicrobiales* type strains is shown, built using the concatenated protein alignments encoded by the perc95_58 gene set (454 genes present in at least 95% of the included strains). The numbers on the nodes indicate the ultra-fast jackknife values using a 40% resampling rate (top numbers) and the SH-aLRT support values (bottom numbers), both calculated from 1000 replicates. Only values below 100 are shown. The scale bar represents the average number of amino acid substitutions per site. To the right of the phylogeny is the current family assignments of each of the 58 strains, followed to the right by the proposed family assignments of each strain.

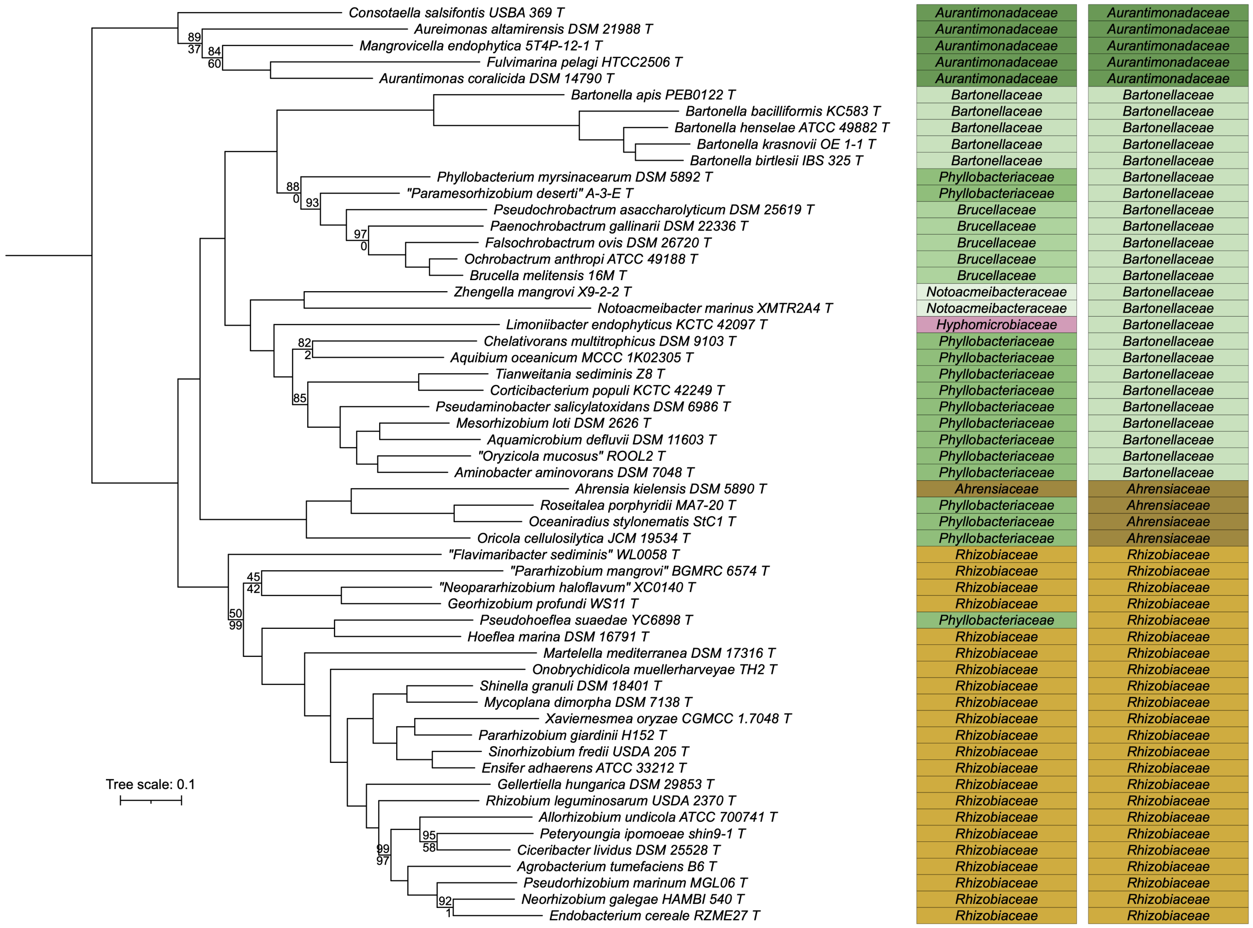

**Figure S11. Phylogenetic analysis of the family *Bartonellaceae* and related families.** On the left, an unrooted maximum likelihood phylogeny of 56 *Hyphomicrobiales* type strains is shown, built using the concatenated protein alignments encoded by the core_56 gene set (178 genes present in 100% of the included strains). The numbers on the nodes indicate the ultra-fast jackknife values using a 40% resampling rate (top numbers) and the SH-aLRT support values (bottom numbers), both calculated from 1000 replicates. Only values below 100 are shown. The scale bar represents the average number of amino acid substitutions per site. To the right of the phylogeny is the current family assignments of each of the 58 strains, followed to the right by the proposed family assignments of each strain.

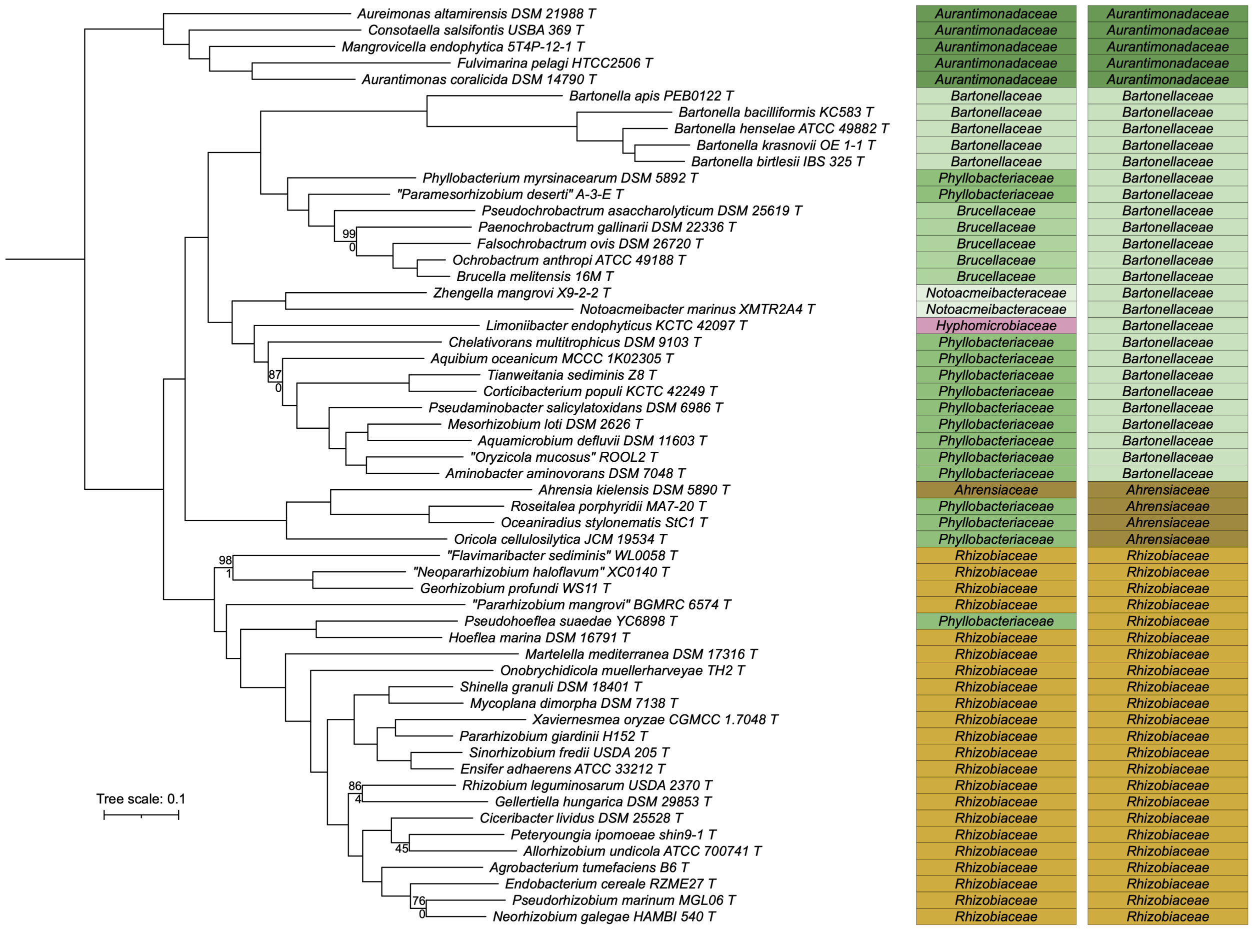

**Figure S12. Phylogenetic analysis of the family *Bartonellaceae* and related families.** On the left, an unrooted maximum likelihood phylogeny of 56 *Hyphomicrobiales* type strains is shown, built using the concatenated protein alignments encoded by the perc95_56 gene set (497 genes present in at least 95% of the included strains). The numbers on the nodes indicate the ultra-fast jackknife values using a 40% resampling rate (top numbers) and the SH-aLRT support values (bottom numbers), both calculated from 1000 replicates. Only values below 100 are shown. The scale bar represents the average number of amino acid substitutions per site. To the right of the phylogeny is the current family assignments of each of the 58 strains, followed to the right by the proposed family assignments of each strain.

### DATASET LEGENDS

**Dataset S1.** Metadata for the 138 *Hyphomicrobiales* strains used in the primary analyses reported in this study, including ftp links to download the genomes.

**Dataset S2.** Metadata for the 5 *Hyphomicrobiales* strains used in this study, including ftp links to download the genomes.

**Dataset S3.** Metadata for the 53 *Hyphomicrobiales* strains used in the secondary analyses of the *Bartonellaceae* and related families, including ftp links to download the genomes.

**Dataset S4.** List of organisms for which the 16S rRNA gene sequences were downloaded from LPSN rather than extracted from the whole genome sequences, including the GenBank accession number of each sequence.

The following datasets are only available through FigShare at:
figshare.com/articles/online_resource/diCenzo_et_al_2023_Hyphomicrobiales_taxonomy/24417334

**Dataset S5.** A matrix of all pairwise cpAAI values calculated from the proteins encoded by the core_143 gene set.

**Dataset S6.** A matrix of all pairwise wpAAI values calculated using EzAAI.

**Dataset S7.** A matrix of all pairwise cpAAI values calculated from the proteins encoded by the core_138 gene set.

**Dataset S8.** The phylogeny of Figure 1 provided in Newick format.

**Dataset S9.** The phylogeny of Figure S1 and S3 provided in Newick format.

**Dataset S10.** The phylogeny of Figure S2 provided in Newick format.

**Dataset S11.** The phylogeny of Figure S4 provided in Newick format.

**Dataset S12.** The phylogeny of Figure S7 provided in Newick format.

**Dataset S13.** The phylogeny of Figure S8 provided in Newick format.

**Dataset S14.** The phylogeny of Figure S9 provided in Newick format.

**Dataset S15.** The phylogeny of Figure S10 provided in Newick format.

**Dataset S16.** The phylogeny of Figure S11 provided in Newick format.

**Dataset S17.** The phylogeny of Figure S12 provided in Newick format.
